# TLR7-agonist and antineoplastic MEK1/2-inhibitor combination unlocks interferon responses from macrophages

**DOI:** 10.1101/615609

**Authors:** Lei Yang, Jeak Ling Ding

## Abstract

Type I interferons are a family of pleiotropic cytokines that exert anti-tumor actions directly on tumor cells and indirectly on the tumor immune microenvironment (TIME). Hitherto, therapeutic strategies aiming to garner the efficacies of interferon responses are still limited. Here we show a novel strategy that elicits an interferon signature response while targeting both tumor cells using antineoplastic mitogen-activated protein kinase (MAPK) kinase 1/2 (MEK1/2) inhibitor and the TIME using toll-like receptor 7 (TLR7)-based immune adjuvant. The combination of MEK1/2 inhibitor and TLR7 agonist unlocked an interferon signature response unexpectedly in macrophages, which was otherwise tightly constrained by TLR7 agonist alone. Deficiency of interferon regulatory factor 1 (*Irf1*) completely abrogated the responses and prevented the reprogramming of activated macrophages, subduing them in an immunosuppressive state. In a murine melanoma model, combination therapy with TLR7 agonist and MEK1/2 inhibitor synergistically extended survival in wild-type but not *Irf1*-deficient mice. Specifically, we identified interferon response genes as favorable prognosis markers for cutaneous melanoma patients. Our findings demonstrate a novel strategy for combination therapy that targets both tumor cells and the immunosuppressive TIME through additive effects of monotherapies and synergistic interferon responses.

## Introduction

Efficacies of anti-cancer therapies are influenced by the specific nature of TIME and the heterogeneous cancer cells that harbor various oncogenic mutations with distinct sensitivities to treatment (1). Numerous molecular drivers of tumorigenesis have been identified, which ultimately converge to major signaling axis like the RAS-RAF-MEK1/2 pathway that have been found in more than 30% of human cancers (2). Although inhibitors targeting the above pathway have generated exciting outcomes, their antineoplastic effects are often compromised by cancer-cell autonomous resistance (3). Intriguingly, a durable anti-tumor immunity requires activation of innate immunity for antigen-presentation, induction of cancer-specific cytotoxic T lymphocytes (CTLs) and constraint of the immunosuppressive TIME (1). The infiltrations of immunosuppressive immune cells like tumor-associated macrophages (TAMs) and regulatory T (Treg) cells are commonly linked to the secretion of inhibitory molecules and poor prognosis, which renders them appealing therapeutic targets to restore the anti-tumorigenic TIME (4, 5). Hence, it is suggested that cancer patients will most likely benefit from therapeutic interventions with both antineoplastic effects on tumor cells and immunostimulatory effects on the TIME (6).

Interferons are known to exert multifaceted anti-tumor actions directly on cancer cells and indirectly on the TIME (7, 8). There are in total three distinct families of interferons, among which the type I interferons possess both antineoplastic and immunomodulatory effects (7). Upon binding to corresponding receptors, all interferons are known to activate the Janus kinase/signal transducers and activators of transcription (JAK/STAT) pathway and subsequently, transcription of diverse anti-tumor interferon-stimulated genes (8). In particular, the antineoplastic effects are mediated by promoting cancer cell apoptosis, inhibiting tumor cell proliferation or by inducing tumor neoantigens, secondary mediators, and major histocompatibility complex I (MHC I) (7, 8). The immunomodulatory actions determine the maturation of dendritic cells (DCs), activation of natural killer (NK) cells, cytotoxicity and survival of cluster of differentiation 8 (CD8^+^) CTLs, antibody response in B cells, and suppressive roles of TAMs and Treg cells (7, 8). These findings have revitalized the earlier notion that type I interferons are the “magic bullet” to cure cancers.

Although polyethylene glycol-conjugated interferon α2b was approved in 2011 for the adjuvant treatment of melanoma patients, systemic administration of type I interferons is marred by dose-limiting side effects like flu-like symptoms and depression (7, 8). Follow-up attempts managed to deliver recombinant type I interferons specifically into the TIME using interferon-conjugated cancer cell-specific antibodies (9) or interferon overexpression in mesenchymal stem cells (10) and monocytes (11). In general, type I interferons are induced by pathogen recognition receptors upon encounter of exogenous or endogenous nucleic acids like double-stranded RNA, single-stranded RNA and CpG DNA (7, 8). Therefore, alternative anti-cancer strategies using pathogen recognition receptor agonists were also exploited to produce type I interferons in plasmacytoid DCs (pDCs) using poly(A:U) (12) and CpG DNA adjuvant (13), TLR7 agonist imiquimod (14) and RNA-lipoplexes (15), or in other immune cells like macrophages using STING agonist (16), Bacillus Calmette–Guérin (17) and poly(I:C) (18). Interestingly, the anti-tumor effects of ionizing radiation (19) and the chemotherapy drug, anthracyclines (20), were also shown to be mediated by cancer cell-autonomous interferon responses, which represents an alternative intervention strategy. However, the above-mentioned therapeutic strategies target mainly the interferon-mediated immunity but not the molecular drivers of tumorigenesis. Hence, alternative intervention strategy is needed to target both cancer cell proliferation and the TIME to mount more effective anti-tumor interferon responses (7).

Downstream of the well-known *RAS* and *RAF* oncogenes, MEK1/2- extracellular signal-regulated kinase (ERK) MAPK pathway has been suggested as one of the most druggable pathways (2), having received four FDA approvals since 2011. Notably for immune responses, the MEK1/2 pathway is explicitly activated by the tumor progression locus 2 (Tpl2, also known as COT or MAP3K8) rather than the RAS and RAF kinases (21). Upon TLR4 and TLR9 activation, *Tpl2* deficiency led to increased expression of interferon-β (Ifn-β, type I interferon) and proinflammatory cytokines like interleukin 12 (IL-12) in macrophages but not in pDCs (22, 23). Our previous findings (24, 25) identified a sustained suppression of TLR3-induced IRF1 in macrophages possibly by the TLR7-MEK1/2 pathway. Given that IRF1 is essential for interferon production (26) and signaling (27, 28), the above suppression may also constrain the interferon-producing capabilities of TLR7-activated macrophages as previously suggested (22, 23). Therefore, we investigated whether the suppression mechanism may be exploited as a novel strategy to elicit interferon responses while maintaining the immunostimulatory roles of TLR7 agonist (14, 29) and the antineoplastic roles of MEK1/2 inhibitors (2, 21).

Here we found, for the first time, that an interferon signature response was unlocked through the synergy between MEK1/2 inhibitor and TLR7 agonist. We elucidated essential roles of the NF-κB-IRF1 signaling axis in unlocking the interferon responses and in reprogramming of macrophages from immunosuppressive to immunostimulatory phenotype. The anti-tumor efficacy of the interferon responses unlocked by the combination treatment strategy was fully substantiated by a murine melanoma model *in vivo*. Our findings demonstrated a novel combination therapy that may be exploited to target the progression of *RAS*/*RAF*-mutated tumor cells, and the transition of TIME from immunosuppressive to immunostimulatory state for a robust and durable anti-tumor immunity.

## Results

### TLR7 activation constrains itself and other TLRs from inducing interferon response genes in macrophages

TLR7 activation in macrophages suppresses IRF1 during TLR crosstalk possibly through the MEK1/2-ERK MAPK pathway (24, 25). As IRF1 is essential for interferon production (26) and responses (27), we postulated that interferon signaling may be similarly altered in macrophages. To test this hypothesis, we stimulated bone-marrow derived macrophages (BMDMs) using poly(I:C) (double-stranded RNA analogue, TLR3 agonist) (24, 25) and/or R848 (resiquimod, TLR7/8 agonist) (24, 25). Single stimulation with R848 did not induce high expression of all six well-known interferon response genes including *Irf1*, *Ifn-β*, *Gbp4* (guanylate-binding protein 4), *Ifit1* (interferon-induced protein with tetratricopeptide repeats 1), *Ifit2* and *Ifit3* (**Fig. 1a**). Consistently, poly(I:C)-induced gene expression was significantly (*P*<0.05) reduced by the addition of R848 (**Fig. 1a**). Similarly, addition of R848 into LPS (lipopolysaccharides, TLR4 agonist)-stimulated macrophages significantly (*P*<0.05) reduced the induction of *Irf1*, *Ifn-β*, *Ifit2* and *Ifit3* (**Fig. 1b**). As TLR3 and TLR4 activate distinct signaling pathways (30), these findings imply a broad-acting suppression mediated by TLR7. At the protein level, TLR7 activation suppressed poly(I:C)- and LPS-induced IRF1 and STAT1 (signal transducer and activator of transcription 1) (**Fig. 1c** and **1d**). Given that STAT1 is an indispensable regulator of interferon signaling (30), the above findings suggest the suppression of both interferon expression and signaling. However, the suppression of IRF1 and STAT1 did not occur in BMDM generated from the myeloid differentiation primary response 88-deficient mice (*Myd88^-/-^*; an indispensable adaptor for TLR7) (**Supplementary Fig. 1a** and **1b**), implying the direct involvement of a TLR7-specifc mechanism.

**Figure 1.**
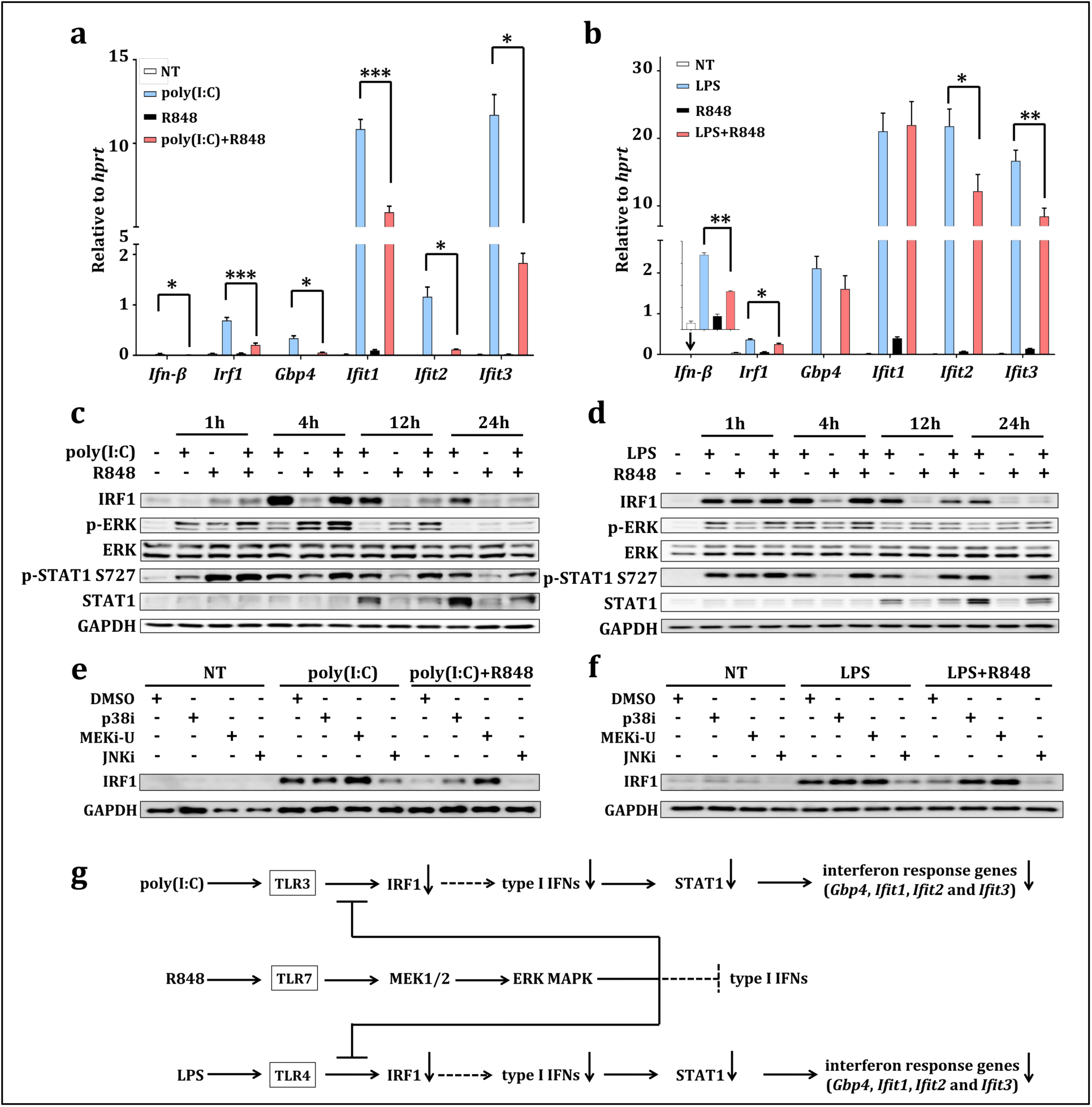
TLR7 stimulation constrains expression of interferon response genes during TLR crosstalk in macrophages. (**a**, **b**), mRNA expression of six interferon response genes (*Ifn-β, Irf1, Gbp4, Ifit1, Ifit2* and *Ifit3*) in BMDM stimulated as indicated for twelve hours. (**c**, **d**) Immunoblot analysis of IRF1, total and p-ERK, total and p-STAT1 in BMDM stimulated for indicated time intervals. (**e**, **f**) Immunoblot analysis of IRF1 in stimulated BMDM. Cells were stimulated with TLR3 agonist poly(I:C) and TLR7 agonist R848 (**e**), or with TLR4 agonist LPS and R848 (**f**) for twelve hours in the presence or absence of indicated MAPK inhibitors. (**g**) schematic illustration of TLR7-specific suppression on TLR3- and TLR4- induced IRF1 and interferon responses. *0.01<*P*<0.05, **0.001<*P*<0.01, \*\*\**P*<0.001 (unpaired Welch’s t-test). Data are presented as mean + *s.e.m.* (**a**-**b**) or one representative of four independent experiments (**c**-**f**). See also **Supplementary Fig. 1**.

MAPKs are known to play dual opposing roles, mediating both the activation and suppression of proinflammatory cytokines (21, 31, 32). In line with our previous findings (24), we identified an inverse relationship between TLR7-mediated suppression and the sustained phosphorylation of ERK MAPK (p-ERK) (**Fig. 1c** and **1d**). Thus, we postulated that MAPKs may be involved in the suppression of interferon response genes. Target-specific inhibition of MEK1/2-ERK MAPK pathway by MEK1/2 inhibitor (MEKi-U) (33), but not inhibition of JNK MAPK (JNKi) (24, 25), rescued both poly(I:C)- and LPS-induced IRF1 production (**Fig. 1e** and **1f**). Although elevated IRF1 production and STAT1 phosphorylation were also observed during the inhibition of p38 MAPK (p38i) (24, 25), no sustained activation of p38 MAPK was associated with the above suppression (**Fig. 1e** and **1f**) as previously suggested (24). Therefore, MEK1/2 pathway, rather than the p38 or JNK MAPK pathways, is responsible for the TLR7-mediated suppression. As the mRNA (**Fig. 1a** and **1b**) and protein (**Fig. 1c** and **1d**) levels of IRF1 showed consistent profiles of expression, the suppression may be mediated by transcriptional or post-transcriptional regulation via mRNA degradation. However, when the *de novo* mRNA synthesis was inhibited, mRNA degradation of both *Irf1* and *Ifn-β* were unaltered by MEKi-U treatment (**Supplementary Fig. 1c**). Hence, the suppression is likely mediated at the transcriptional level. Collectively, we identified a TLR7-specific suppression, which causes incompetency of interferon responses (**Fig. 1g**) in TLR7-activated macrophages via the MEK1/2 pathway.

### MEK1/2 inhibitor synergizes with TLR7 agonist to unlock an interferon signature response

Both the interferons and IRF1 are fundamental immune modulators that activate an array of interferon-stimulated genes with diverse essential functions (7, 26-28, 30). Remarkably, TLR7 agonist-based adjuvants are known to mount interferon responses via pDCs (14), through which multiple cell types are activated in various cancers (7, 29, 30). Hence, subversion of the abovementioned TLR7-specific suppression (**Fig. 1g**) in macrophages may represent a novel strategy to fully elicit the interferon responses.

To determine whether the above TLR7-suppression (**Fig. 1e** and **1f**) is specific for the interferon response genes, we performed a whole transcriptome analysis. Primary macrophages were treated with R848 to activate the TLR7 signaling, and with MEKi-U (U0126; IC_50_=72 nM) (33) to inhibit the MEK1/2 pathway (**Supplementary Fig. 2a**). Compared to the dimethyl sulfoxide (DMSO) vehicle control, there were about 32% more differentially expressed genes (DEGs; fold change>1.5; adjusted *P*<0.05) when MEK1/2 pathway was inhibited (777 versus 1024) (**Supplementary Fig. 2b**). Based on the dual regulatory roles of MEK1/2 pathway, common DEGs with larger than 2-fold difference (adjusted *P*<0.05) between the DMSO- and MEKi-U- treated groups were shortlisted into two categories: **Activation** and **Suppression** (**Fig. 2a**). Amongst genes under suppression of the MEK1/2 pathway, 33% (39 out of 117) and 24% (28 out of 117) were able to interact with STAT1 (**Fig. 2a**, **purple** color) and IRF1 (**Fig. 2a**, **orange** color), respectively. Therefore, STAT1 and IRF1 are possibly involved in the regulation of these genes. We found that the most highly enriched gene sets activated by the MEK1/2 pathway were those related to the negative regulation of myeloid cell activation and response (**Fig. 2b**). In contrast, the suppressed genes were highly enriched in gene sets exhibiting an “interferon signature response” (**Fig. 2b**), indicating that subversion of the above suppression is indeed a promising strategy to unlock macrophage-based interferon responses.

**Figure 2.**
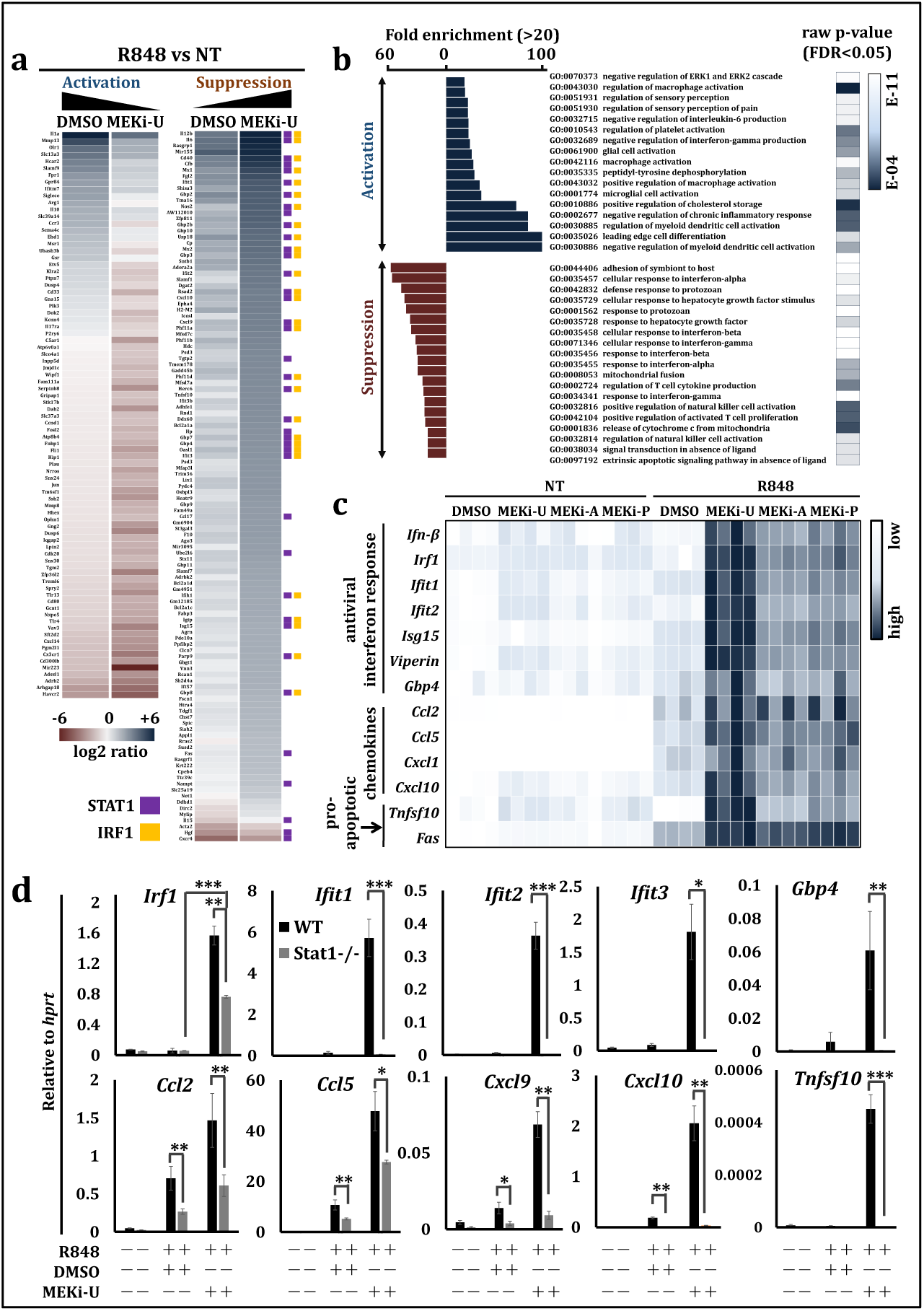
MEK1/2 inhibitor synergizes with TLR7 agonist to unlock an interferon signature response. (**a**) Heat map showing whole transcriptional analysis of differentially expressed genes (DEGs) activated (**Activation**) or suppressed (**Suppression**) by the MEK1/2 pathway. BMDM were treated with TLR7 agonist R848 in the presence or absence of MEK1/2 inhibitor MEKi-U as shown in **Supplementary Fig. 2a**. DEGs interacting with STAT1 and IRF1 were labeled with **purple** and **orange** squares based on STRING PPI test. (**b**) Gene set enrichment analysis using DEGs listed in (**a**). Only gene sets with fold change (raw *P*-value<10^−4^ with FDR<0.05) higher than twenty folds were presented. (**c**) Heat map showing mRNA expression of seven interferon response genes (*Ifn-β*, *Irf1*, *Ifit1*, *Ifit2*, *Isg15*, *Viperin* and *Gbp4*), four chemokines (*Ccl2*, *Ccl5*, *Cxcl1*, *Cxcl10*) and two pro-apoptotic genes (*Tnfsf10* and *Fas*). BMDM were activated by R848 for six hours in the presence or absence of three MEK1/2 inhibitors (MEKi-U, MEKi-A and MEKi-P). Scale bars are presented as a range from low to high expression relative to each gene. (**d**) mRNA expression of five interferon response genes (*Irf1, Ifit1*, *Ifit2, Ifit3* and *Gbp4*), four chemokines (*Ccl2*, *Ccl5, Cxcl9* and *Cxcl10*) and one pro-apoptotic gene (*Tnfsf10*) in BMDM generated from WT and *Stat1*^-/-^ mice. Cells were activated similarly as in (**a**). *0.01<*P*<0.05, **0.001<*P*<0.01, \*\*\**P*<0.001 (unpaired Welch’s t-test or two-way ANOVA with Tukey’s correction). Data are presented as relative expression (**c**) or mean ± S.D. (**d**) of four independent experiments. See also **Supplementary Fig. 2**.

To verify the above interferon signature response, we used three widely tested MEK1/2 inhibitors MEKi-U, MEKi-A (AZD6244, Selumetinib; IC_50_=14 nM) (2, 34) and MEKi-P (PD0325901; IC_50_=0.33 nM) (2) to inhibit the MEK1/2 pathway and then, measured the expression of interferon response genes. Consistently, thirteen genes showed elevated expression during treatment with inhibitors (**Fig. 2c**), including seven interferon response genes (*Ifn-β*, *Irf1*, *Ifit1*, *Ifit2*, *Isg15*, *Viperin* (virus inhibitory protein, endoplasmic reticulum-associated, interferon-inducible), and *Gbp4*), four chemokines (*Ccl2* (C-C motif chemokine ligand 2), *Ccl5*, *Cxcl1* (C-X-C motif chemokine ligand 1) and *Cxcl10*), and two pro-apoptosis genes (*Tnfsf10* (TNF superfamily member 10), and *Fas* (Fas cell surface death receptor)). In contrast, expression of the pro-inflammatory cytokine gene, interleukin 1α (*Il-1α*) was significantly (*P*<0.01) reduced (**Supplementary Fig. 2c**), consistent with our data and previous report (22). Notably, the p38 MAPK was not involved in the unlocked interferon signature response as its inhibitor only mildly increased the expression of interferon response genes (**Supplementary Fig. 2d**). These findings provide insights into the hitherto uncharacterized incompetency of macrophages to induce interferons upon TLR7 activation, which may be further exploited as a novel strategy to elicit interferon responses.

### Enhanced type I interferon signaling leads to unlocked interferon signature response

To investigate how the interferon signature response is unlocked, we tested the direct roles of interferons. As the MEK1/2 inhibitor did not alter *Ifn-α* expression in TLR7-activated macrophages, it is unlikely for IFN-α to be involved (**Supplementary Fig. 2e**). Using antigen-specific antibodies, we blocked the cell surface receptors for type I interferons (IFNAR, receptor for IFN-α and IFN-β) and type II interferon (IFNGR, receptor for IFN-γ). During treatment with MEK1/2 inhibitor and TLR7 agonist, STAT1 activation was enhanced in immunoglobulin G (IgG) control group but was reduced only after IFNAR neutralization (**Supplementary Fig. 2f**), indicating the sole involvement of IFN-β. Furthermore, as compared to those from wild-type (WT) mice, BMDM generated from the *Stat1*^-/-^ mice expressed significantly (*P*<0.05) lower or even abrogated levels of five interferon response genes (*Irf1*, *Ifit1*, *Ifit2, Ifit3* and *Gbp4*), four chemokines (*Ccl2, Ccl5, Cxcl9* and *Cxcl10*) and pro-apoptosis gene *Tnfsf10* (**Fig. 2d**). However, pro-apoptotic gene *Fas* and chemokine *Cxcl1* were not affected (**Supplementary Fig. 2g**). Such discrepancies in mediating the synergistic interferon signature response suggest the essential but selective roles of IFN-β-STAT1 signaling pathway and the possible involvement of additional signaling pathways.

### Unlocked IRF1 mediates an autocrine IRF1-Interferon-STAT1-IRF1 signaling axis

In macrophages, interferons are known to activate an autocrine amplification loop either directly via the STAT1-STAT2-IRF9 complex and STAT1-STAT1 homodimers, or indirectly via transcription factors like IRF1 (27). Notably, IRF1 also mediates the interferon production (26, 28). Therefore, the interplay between IRF1 and the unlocked interferon signature response suggests that IRF1 may function both downstream and upstream of the interferon-STAT1 signaling.

We first tested its function downstream of the interferon signaling using recombinant IFN-β and IFN-γ (27). Only IFN-β- but not IFN-γ- induced IRF1 production was suppressed by the addition of TLR7 agonist R848 (**Supplementary Fig. 3a**). Similarly, antibody-based neutralization of IFNAR but not of IFNGR reduced the unlocked IRF1 (**Supplementary Fig. 3b**). These findings revealed a TLR7-specific suppression on both signaling and expression of type I interferon. We then reconstituted STAT1 protein in macrophages lacking endogenous *Stat1* gene (35) and found a slight increase of *Irf1* expression (**Supplementary Fig. 3c** and **3d**). Remarkably, this increased expression was not comparable to the level unlocked by the MEK1/2 inhibitor treatment (**Fig. 2c**), suggesting a partial role of STAT1 in regulating IRF1 expression. Therefore, these findings revealed the partial involvement of IRF1 downstream of the interferon-STAT1 signaling pathway.

We next determined whether IRF1 functions as an upstream regulator. TLR7 activation by R848 induced a transient expression of *Irf1* shortly after stimulation but not at later time points (**Supplementary Fig. 3e**). Such kinetics is associated with the incompetency of interferon production in TLR7-activated macrophages (**Fig. 2c**). We then investigated whether IRF1 mediates the interferon signature response. Compared to WT counterparts, all interferon response genes tested were significantly (*P*<0.05) less expressed in *Irf1*-deficient (*Irf1*^-/-^) BMDMs (**Fig. 3a** and **Supplementary Fig. 3f**). We next probed the roles of IRF1 using whole transcriptome analysis (**Supplementary Fig. 3g**). Compared to the common DEGs (adjusted *P*<0.05) that differ by less than 1.5 folds between the WT and *Irf1^-/-^* macrophages (**Fig. 3b**), DEGs with higher fold changes (FC) were enriched in the interferon response-related signatures (**Fig. 3c** and **3d**). Remarkably, these enriched gene sets largely overlapped with those shown above for the interferon signature response (**Fig. 2b**), supporting the role of IRF1 as an upstream regulator. Collectively, IRF1 is a *bona fide* mediator of the autocrine IRF1-Interferon-STAT1-IRF1 signaling axis.

**Figure 3.**
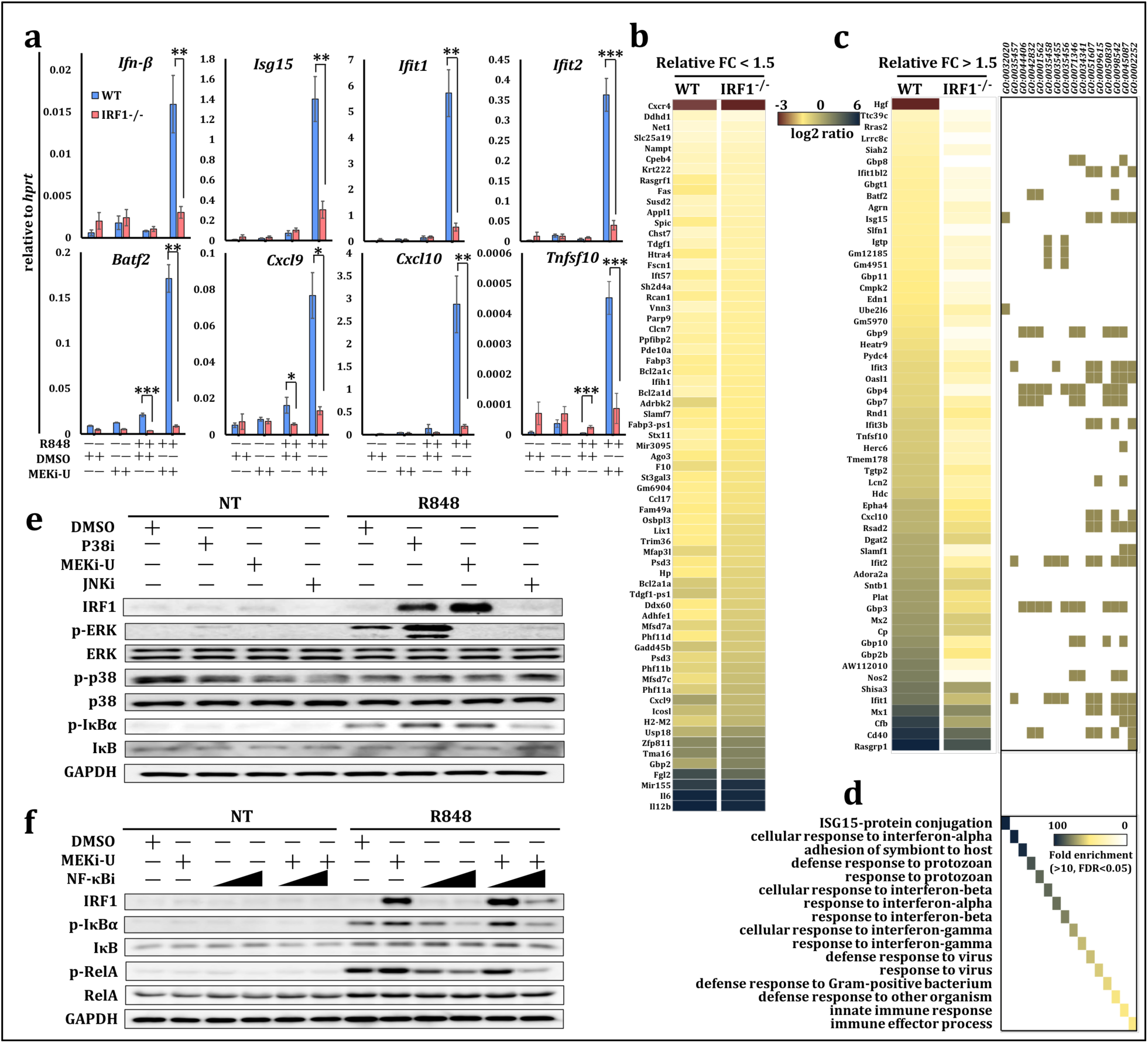
Unlocked interferon signature response is IRF1-dependent. (**a**) mRNA expression of four interferon response genes (*Ifn-β*, *Isg15*, *Ifit1* and *Ifit2*), one master transcription factor (*Batf2*), two chemokines (*Cxcl9* and *Cxcl10*) and one pro-apoptosis gene (*Tnfsf10*) in BMDM generated from WT and *Irf1*^-/-^ mice. Cells were stimulation with TLR7 agonist R848 for six hours in the presence or absence of MEK1/2 inhibitor MEKi-U. (**b**, **c**) Heat map showing DEGs from whole transcriptome analysis using both WT and *Irf1*^-/-^ BMDM. Cells were treated in the same way as in (**a**). Genes were divided into two groups with less (**b**) or no less (**c**) than 1.5-fold differences between WT and *Irf1*^-/-^ BMDM. FC, fold change. (**d**) Gene set enrichment analysis of DEGs listed in (**c**). (**e**) Immunoblot analysis of IRF1, total and p-ERK, total and p-p38, total and p-IκB. BMDM were activated by R848 for twelve hours in the presence or absence of MEKi-U, p38 MAPK inhibitor p38i, or JNK MAPK inhibitor JNKi. (**f**) Immunoblot analysis of IRF1, total and p-IκB, total and p-RelA (p65). BMDM were stimulated with R848 for twelve hours in the presence or absence of MEKi-U and NF-κB inhibitor NF-κBi (1 μM and 5 μM). *0.01<*P*<0.05, **0.001<*P*<0.01, \*\*\**P*<0.001 (unpaired Welch’s t-test). Data are presented as mean ± S.D. (**a**) or one representative of four independent experiments (**e**, **f**). See also **Supplementary Fig. 3.**

### IRF1 is unlocked through the activation of NF-κB pathway

The above partial dependence of *Irf1* expression on STAT1 (**Fig. 2d**) suggests contributions from other IRF1-inducing signaling pathways such as the NF-κB pathway. Although functions of the NF-κB pathway are generally implicated in inflammatory responses (30), its inhibition was found to restore both inflammatory and antiviral responses (36). Therefore, we postulated that NF-κB pathway mediates the unlocked IRF1 activity. Indeed, unlike the total NF-κB inhibitor α (IκBα), the phosphorylated form (p-IκBα) was largely increased following combination treatment (**Fig. 3e**). To confirm this association with the unlocked IRF1, we treated macrophages with both MEKi-U and NF-κB inhibitor (NF-κBi). Consistently, MEKi-U largely elevated IRF1 production when administered together with TLR7 agonist (**Fig. 3f**). However, the effect of MEK1/2 inhibitor on IRF1 production was abolished when administered together with NF-κBi (**Fig. 3f**), highlighting the vital role of the NF-κB pathway. The increase in p-IκBα augmented the activities of NF-κB pathway as manifested by a rise in the p-RelA, a central effector of NF-κB pathway (**Fig. 3f**). We then concluded that NF-κB pathway is responsible for the unlocked IRF1 and interferon signature response.

### Combination treatment reprograms macrophages towards a predominantly M1-like phenotype through IRF1-Interferon-STAT1 pathway

Macrophages are highly versatile and can easily undergo reprogramming in response to the rapidly changing environmental cues (5, 37). Classically activated M1 macrophages are known to be immunostimulatory during immunotherapy, whereas alternatively activated M2 macrophages exert immunosuppressive effects in diverse established tumors (5). Our above findings identified significant increases of *Il-12b* and *Il-6* in macrophages under both MEK1/2 inhibitor and TLR7 agonist treatment (**Supplementary Fig. 2d**). As both cytokines are commonly found in M1-like macrophages (36), we hypothesized that the combination treatment may drive the phenotypic change of stimulated macrophages via IRF1-mediated responses or *vice versa*.

To elucidate the causative links, we tested the phenotypic changes of activated macrophages. We found a mixed phenotype after treatment with TLR7 agonist R848. In particular, there were increased expression of both M1 phenotype markers including nitric oxide synthase 2 (*Nos2*), *Il-12b* and tumor necrosis factor α (*Tnf-α*) (**Fig. 4a**), and M2 phenotype marker arginase 1 (*Arg1*) (**Fig. 4b**). Similar results were observed when examining M1 marker MHC II (major histocompatibility complex II) (**Fig. 4c**) and M2 markers CD80 (high-affinity ligand for immune checkpoint CD152/CTLA-4), CD163 (**Fig. 4d**) and the mannose receptor C-type 1 (*Mrc1*, also known as CD206) (**Fig. 4b**). In contrast, MEK1/2 inhibitor modulated only two phenotype markers, including MHC II (**Fig. 4c**) and CD80 (**Fig. 4d**), whereas other markers remained unchanged (**Fig. 4a-4d**). However, macrophages during combination treatment predominantly exhibited M1-like phenotype with elevated M1 (**Fig. 4a** and **4c**) and decreased M2 phenotype markers (**Fig. 4b**, **4d** and **4e**).

**Figure 4.**
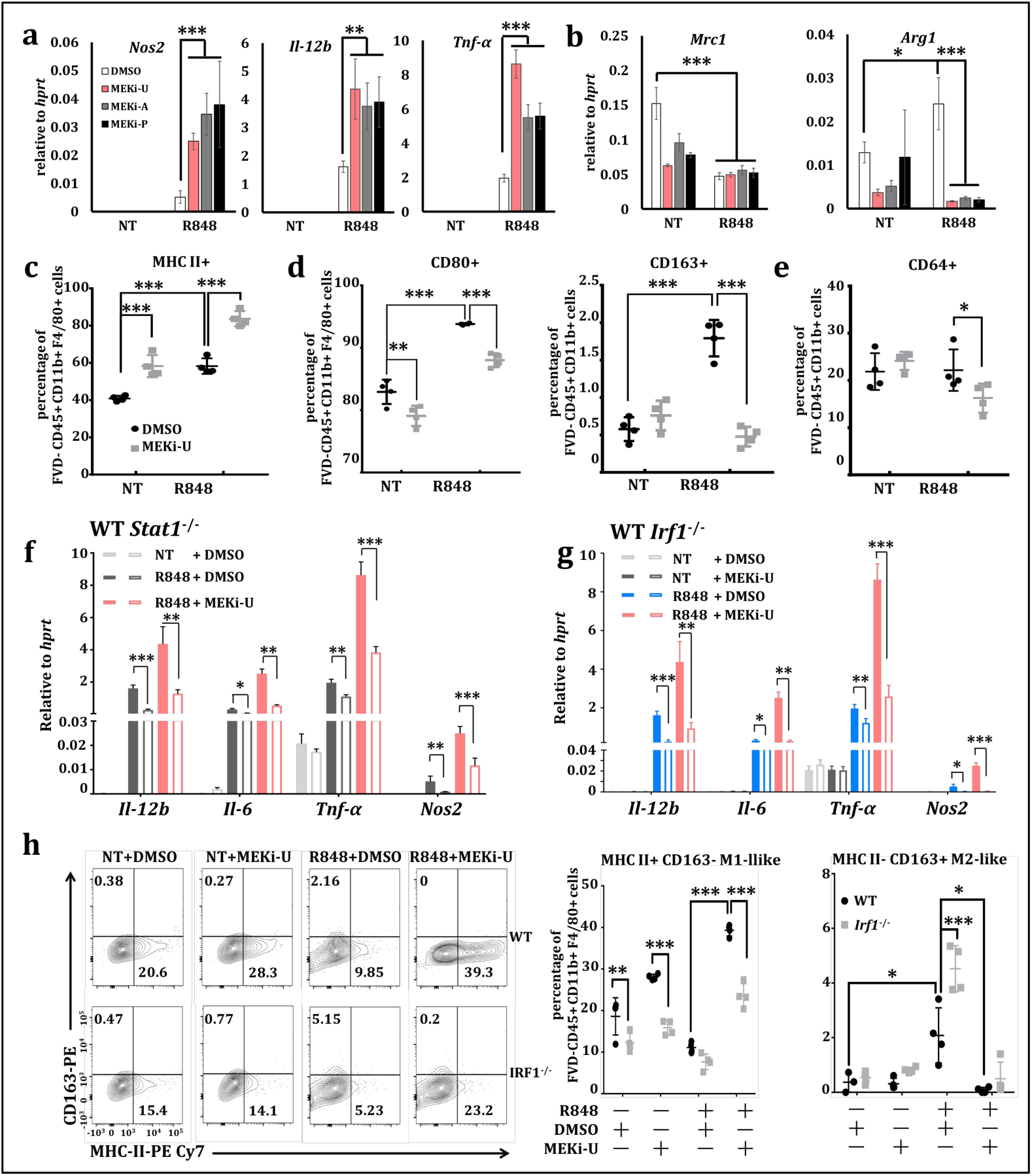
Combination treatment with MEK1/2 inhibitor and TLR7 agonist reprograms macrophages towards a predominantly M1-like phenotype through IRF1. (**a**, **b**) mRNA expression of three M1 phenotype genes (*Nos2*, *Il-12b* and *Tnf-α*) (**a**) and two M2 phenotype genes (*Mrc1* and *Arg1*) (**b**). BMDM were activated via TLR7 agonist R848 for six hours in the presence or absence of three MEK1/2 inhibitors. (**c**, **d**, **e**) Surface expression of M1 phenotype marker MHC-II (**c**), M2 phenotype markers CD80 and CD163 (**d**) and phagocytosis marker CD64 (**e**) within the indicated cell populations. BMDM were stimulated with R848 for twenty-four hours in the presence or absence of MEKi-U. Cells were then harvested and assayed for flow cytometry analysis. (**f, g**), mRNA expression of four M1 phenotype genes (*Il-12b, Il-6, Tnf-α* and *Nos2*) in BMDM generated from WT and *Stat1*^-/-^ mice (**f**) or from WT and *Irf1*^-/-^ mice (**g**). Cells were stimulated with R848 for six hours in the presence or absence of MEKi-U. (**h**) Percentages of MHC II^+^ CD163^-^ M1 and MHC II^-^ CD163^+^ M2 phenotype markers shown by representative flow cytometry analysis (left) and corresponding quantifications (right) in BMDM generated from WT and *Irf1*^-/-^ mice. Cells were similarly treated and processed as that in (**c-e**). *0.01<*P*<0.05, **0.001<*P*<0.01, \*\*\**P*<0.001 (one-way ANOVA with Bonferroni’s correction or two-way ANOVA with Tukey’s correction). Data are presented as mean ± S.D. or one representative of four (**a**-**g**) or two (**h**-**k**, n=4 per group) independent experiments. See also **Supplementary Fig. 4.**

To study the direct involvement of IRF1-mediated interferon signature response, macrophage phenotype markers in BMDMs generated from WT, *Stat1*^-/-^ and *Irf1*^-/-^ mice were examined. Deficiencies of both *Stat1* (**Fig. 4f**) and *Irf1* (**Fig. 4g**) led to significantly (*P*<0.05) lower expression of all four M1 phenotype markers (*Il-12b*, *Il-6*, *Tnf-α* and *Nos2*). At the cellular level, *Irf1* deficiency promoted M1-like (MHC II^+^ CD163^-^) macrophages but constrained the M2-like (MHC II^-^ CD163^+^) macrophages (**Fig. 4h**). These findings clearly demonstrated the novel and indispensable roles of IRF1 and STAT1 in reprogramming M2-like macrophages towards an immunostimulatory M1-like phenotype.

### MEK1/2 inhibitor dampens proliferation of naïve and TLR7-activated macrophages

In addition to its roles in the immune responses, MEK1/2 pathway is also known to modulate immune cell proliferation and survival (13, 38, 39). Consistently in our studies, MEK1/2 inhibitor treatment significantly increased cell death of naïve macrophages (*P*<0.001) but did not affect the viabilities (*P*>0.05) of macrophages when combined with R848 (**Supplementary Fig. 4a**). The proliferative capabilities of macrophages were similarly reduced in both naïve and activated states after MEK1/2 inhibitor treatment (**Supplementary Fig. 4b**). As a result of the reduced proliferation capabilities, mature CD11b^+^ and CD68^+^ macrophages were significantly (*P*<0.001) reduced during combination treatment (**Supplementary Fig. 4c**). Collectively, we concluded that the MEK1/2 pathway exerts dual regulatory effects on both macrophage polarization and proliferation.

### IL-10 signaling mediates the TLR7-specific suppression on IRF1-mediated interferon signature response

MEK1/2-ERK MAPK pathway is known to phosphorylate over 400 substrates in a stimulus-specific manner, of which the mitogen- and stress-activated protein kinases 1 and 2 (MSK1/2)-IL-10 pathway was proven to suppress TLR responses (31, 32). During TLR3-TLR7 crosstalk, *Il-10* expression (**Supplementary Fig. 5a**) and STAT3 phosphorylation (indispensable for the IL-10 signaling) (**Supplementary Fig. 5b**) were largely increased. Similar results were also observed during TLR4-TLR7 crosstalk (**Supplementary Fig. 5c**). These findings prompted us to examine the roles of IL-10 signaling in the TLR7-specific suppression.

In TLR7-activated macrophages, the *Il-10* expression level peaked within an hour post-stimulation, which is reciprocal to the suppression of IRF1-mediated interferon signature response (**Supplementary Fig. 5d**). Remarkably, R848 stimulation induced the largest amount of IL-10 protein compared to all other TLR agonists tested, including the poly(I:C), Pam3CSK4 (TLR1/2 agonist) and LPS (**Fig. 5a**). Furthermore, TLR7 activation did not induce *Il-10* during treatment by all three MEK1/2 inhibitors (**Fig. 5b**) but did so during treatment with p38i (**Supplementary Fig. 5e**). This functional discrepancy between MEK1/2 pathway and p38 MAPK pathway explains their distinct roles on the interferon signature response as shown above (**Supplementary Fig. 2d** and **Fig. 3e**). Notably, *Stat1*^-/-^ macrophages displayed significantly (*P*<0.05) reduced *Il-10* regardless of MEK1/2 inhibitor (**Supplementary Fig. 5f**). Hence, besides the role in IRF1-mediated interferon signature response, STAT1 also partially mediates the IL-10 signaling as previously suggested (32, 40).

**Figure 5.**
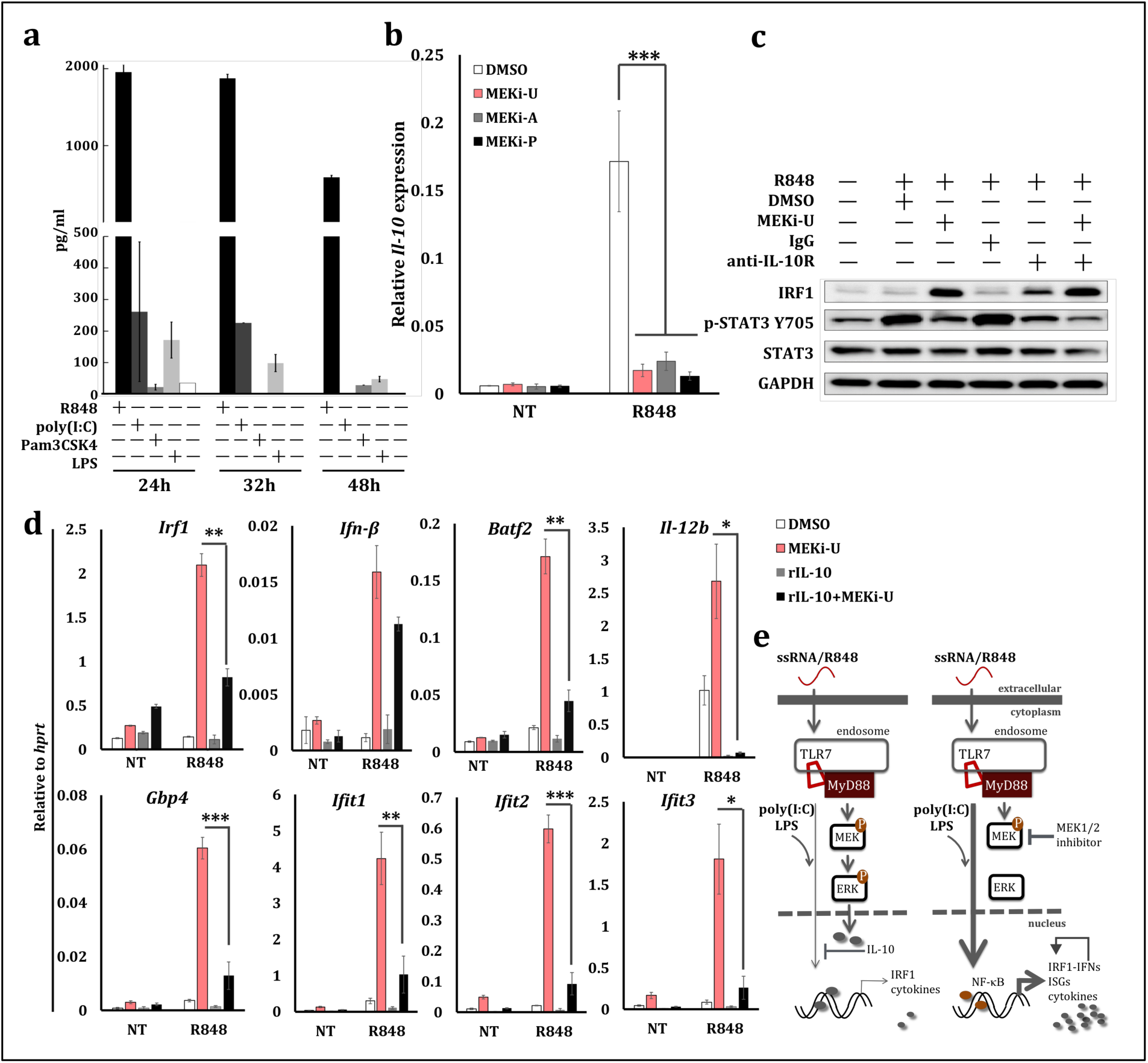
MEK1/2 pathway constrains IRF1-mediated interferon signature response through the IL-10 signaling. (**a**) IL-10 secretion by BMDM stimulated with TLR7 agonist R848, TLR3 agonist poly(I:C), TLR1/2 agonist Pam3CSK4 and TLR4 agonist LPS for indicated time durations. (**b**) *Il-10* mRNA expression in BMDM stimulated with R848 for six hours in the presence or absence of three MEK1/2 inhibitors. (**c**) Immunoblot analysis of IRF1, total and p-STAT3. BMDM were treated with R848 for eight hours in the presence or absence of MEKi-U, anti-IL-10R antibody and IgG isotype control. (**d**) mRNA expression of interferon response genes (*Irf1*, *Ifn-β, Gbp4, Ifit1, Ifit2* and *Ifit3*), transcription factor *Batf2*, and cytokine *Il-12b*. BMDM cells were stimulated with R848 and recombinant IL-10 (20 ng/ml) for six hours in the presence or absence of MEKi-U. (**e**) A schematic illustration on how TLR7-specific MEK1/2-ERK MAPK pathway modulates TLR4 agonist LPS- and TLR3 agonist poly(I:C)-activated interferon signaling. Role of IL-10 pathway in the suppression of NF-κB-IRF1-mediated interferon signature response is highlighted. *0.01<*P*<0.05, **0.001<*P*<0.01, \*\*\**P*<0.001 (unpaired Welch’s t-test or one-way ANOVA with Bonferroni’s correction). Data are presented as mean ± S.D. or one representative of at least three (**a**, **c**) and four (**b**, **d**) independent experiments. See also **Supplementary Fig. 5**.

We next tested the functional roles of IL-10 signaling. During the combination treatment in macrophages, the blockade of IL-10 receptor (IL-10R) unlocked IRF1 production by decreasing the STAT3 phosphorylation (**Fig. 5c**). Similar results were observed during TLR3-TLR7 crosstalk (**Supplementary Fig. 5g**). These findings highlighted the essential roles of IL-10 signaling in the TLR7-specific suppression. Notably, MEK1/2 inhibitor did not alter TLR7-induced *Il-10* expression an hour post-stimulation (**Supplementary Fig. 5h**), which clearly explains the expression kinetics of *Irf1* (**Supplementary Fig. 3e**). Consistent with the previous implications (31, 32), the addition of recombinant IL-10 protein significantly constrained the unlocked expression of interferon response genes by MEK1/2 inhibitor (**Fig. 5d** and **Supplementary Fig. 5i**). Collectively, the above *in vitro* findings revealed a direct involvement of TLR7-MEK1/2-ERK MAPK-IL-10 signaling axis (**Fig. 5e**) in the suppression of NF-κB-IRF1-mediated interferon signature response.

### Combination therapy with MEK1/2 inhibitor and TLR7 agonist improves survival in a murine melanoma model

Our above findings have revealed the potentials of TLR7 agonist and MEK1/2 inhibitor as a novel strategy to elicit IRF1-mediated interferon signature response *in vitro*. We hypothesized that the combination treatment with TLR7 agonist and MEK1/2 inhibitor may exploit the *in vivo* anti-tumor efficacies of type I interferons (7, 8), while fully maintaining the immunostimulatory roles of TLR7 agonist (7, 14, 29) and the antineoplastic roles of MEK1/2 inhibition (2, 21).

Consistent with previous studies (36), we identified high infiltrations of M2-like TAMs (CD80^+^, CD163^+^ and CD206^+^) in a murine model of subcutaneous melanoma (**Supplementary Fig. 6a**). We then treated melanoma-bearing WT mice with monotherapies (either TLR7 agonist or MEK1/2 inhibitor) or combination therapy (both TLR7 agonist and MEK1/2 inhibitor) (**Fig. 6a**). When compared to the vehicle controls, monotherapy with MEK1/2 inhibitor did not alter the survival (*P*=0.3314), possibly due to its selective effects on *RAS*/*RAF*-mutated cancers (41). Hence, the additive effects of monotherapies could be best represented by the impact of TLR7 agonist monotherapy (*P*=0.0003) (**Fig. 6b**). Remarkably, the combination therapy significantly improved the survival (**Fig. 6b**) and delayed the tumor progression (**Fig. 6c**), even in comparison with the additive effects of the two monotherapies (*P*<0.01). The improved efficacy during combination therapy is associated with the unlocked IRF1-mediated interferon signature response, highlighting the well-sustained anti-tumor efficacies of TLR7 agonist and its synergy with MEK1/2 inhibitor.

**Figure 6.**
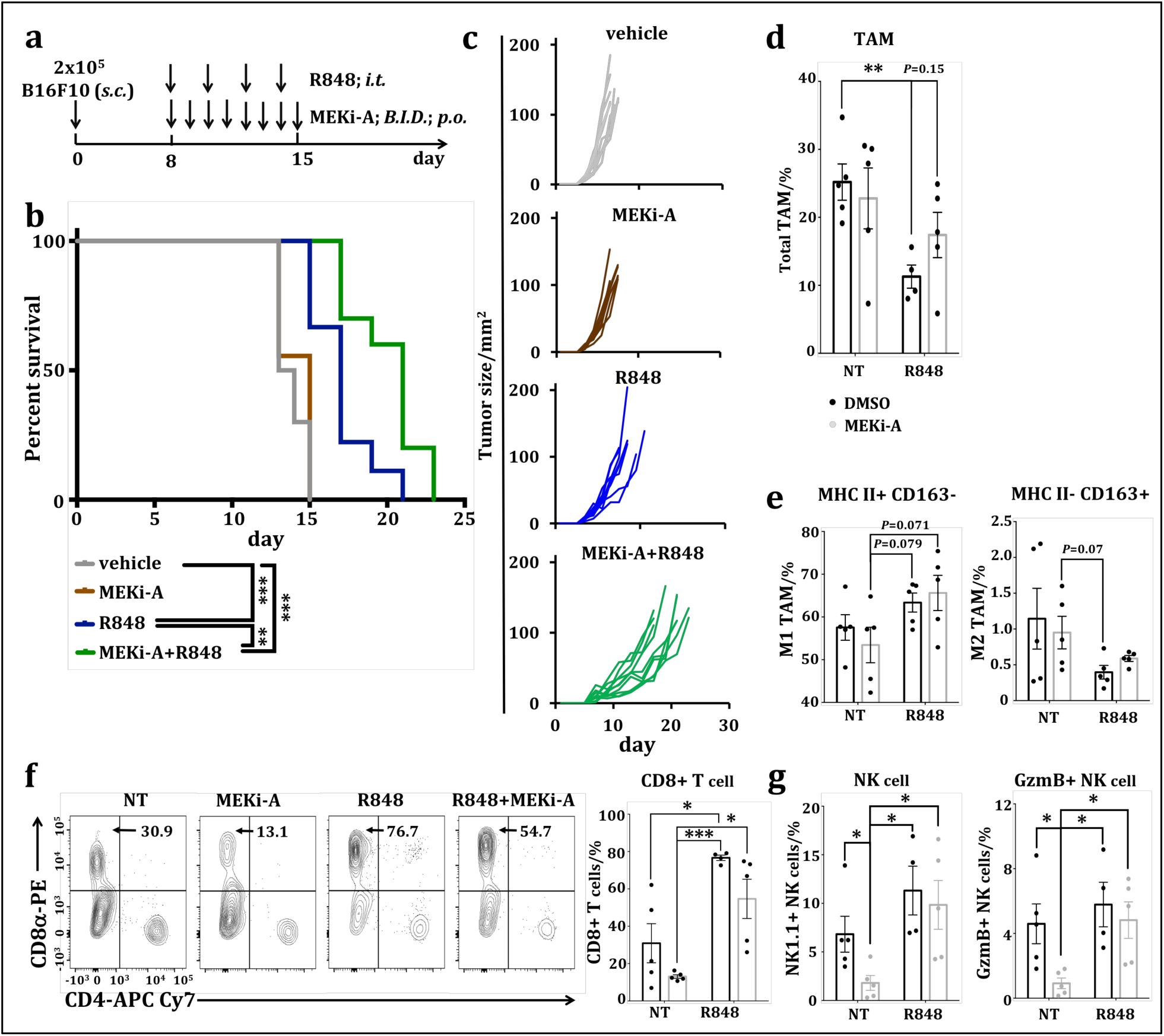
Combination therapy with MEK1/2 inhibitor and TLR7 agonist reduces tumor progression in vivo. (**a**) Treatment scheme of murine B16F10 subcutaneous melanoma model. *s.c.*, subcutaneous. *i.t.*, intratumorally. *B.I.D.*, twice daily. *p.o.*, oral gavage. (**b**) Kaplan–Meier survival curves of WT tumor-bearing mice receiving vehicle (n=9), monotherapy with MEKi-A (n=10), monotherapy with TLR7 agonist R848 (n=10) or combination therapy (MEKi-A and R848; n=10). B16F10 subcutaneous melanoma was established and treated as shown in (**a**). (**c**) individual tumor growth curve of all tumor-bearing mice from (**b**). (**d**, **e, f**, **g**) flow cytometry analysis of B16F10 tumors treated with the scheme shown in (**a**). (**d**, **e**) percentage quantifications of total TAM (FVD^-^ CD45^+^ Ly6G^-^ CD11b^+^ F4/80^+^) in the viable CD45^+^ immune cells (**d**) and MHC II^+^ CD163^-^ M1-like TAMs and MHC II^-^ CD163^+^ M2-like TAMs within the total TAM populations (**e**). (**f**) Percentages of CD4^+^ and CD8^+^ T cells in the FVD^-^ CD45^+^ NK1.1^-^ CD3ε^+^ populations shown by the representative flow cytometry analysis (left) and corresponding quantifications (right). (**g**) percentage quantifications of NK cells (FVD^-^ CD45^+^ CD3ε^-^ NK1.1^+^) in the viable CD45^+^ immune cells and Gzmb+ cells within the NK cell populations. *0.01<*P*<0.05, **0.001<*P*<0.01, \*\*\**P*<0.001 (log-rank test, unpaired Welch’s t-test, or two-way ANOVA with Tukey’s correction). Data are presented as mean ± S.D. or one representative of two (**d**-**g**, n=4 or 5 per group) independent experiments. See also **Supplementary Fig. 6**.

In line with our *in vitro* findings, B16F10 tumor-bearing mice receiving combination therapy have much lower numbers of TAMs (**Fig. 6d**) and MHC II^-^ CD163^+^ M2-like TAMs (**Fig. 6e**), but higher numbers of MHC II^+^ CD163^-^ M1-like TAMs (**Fig. 6e**). However, the pro-tumor immune cells like neutrophils (**Supplementary Fig. 6b**) and Foxp3^+^ Treg cells (**Supplementary Fig. 6c**) remained unchanged, precluding these cells from modulating mice survival. Consistent with previous findings (39), monotherapy with MEK1/2 inhibitor slightly increased the granzyme B^+^ (GzmB^+^) CTLs (**Supplementary Fig. 6d** and **6e**). In contrast, both the TLR7 agonist monotherapy and the combination therapy resulted in significantly (*P*<0.05) higher infiltrations of total CD8^+^ T cells (**Fig. 6f**), total NK cells and GzmB^+^ NK cells (**Fig. 6g** and **Supplementary Fig. 6e**) than monotherapy with MEK1/2 inhibitor. The associations between enhanced infiltrations of tumor-killing effector cells (CD8^+^ T cells, Gzmb^+^ CTLs and NK cells) and enhanced overall survival further substantiate the efficacies of combination therapy. Remarkably, the cellular compositions during monotherapy with TLR7 agonist and combination therapy were similar (**Fig. 6d-6f** and **Supplementary Fig. 6b–6d**) in spite of their differences in mice survival (**Fig. 6b**). This suggests the concurrent involvement of an effector cell infiltration-independent mechanism specifically during the combination therapy.

### Irf1 deficiency abrogates the improved survival during combination therapy

To directly test the involvement of IRF1-medited interferon signature response, we treated both the WT and *Irf1*^-/-^ mice bearing B16F10 tumors with the combination therapy. In line with the increased susceptibility to tumorigenesis of *Irf1*^-/-^ mice (42), we observed significant (*P*<0.001) reduction in mice survival in untreated *Irf1*^-/-^ mice than those of the WT controls (**Supplementary Fig. 6f**). In spite of this, both monotherapy with TLR7 agonist R848 and the combination therapy (R848 and MEK1/2 inhibitor) largely extended the survival of *Irf1*^-/-^ mice, albeit with significantly (*P*<0.001) reduced effects when compared to the WT mice (**Supplementary Fig. 6f**). The compromised efficacy of combination therapy in *Irf1*^-/-^ mice clearly indicates the indispensable anti-tumor role of IRF1. Remarkably, we did not observe any difference in survival (*P*=0.71) in *Irf1*^-/-^ mice receiving monotherapy and combination therapy, suggesting that *Irf1* deficiency abrogated the synergistic interferon signature response observable in WT mice (*P*<0.001) (**Supplementary Fig. 6f**). Altogether, our findings demonstrated a novel strategy that mounted a more superior anti-tumor response than respective monotherapies while maintaining the anti-tumor actions of TLR7 agonist in *RAS*/*RAF* wild-type tumors.

### The interferon signature response indicates favorable prognosis in melanoma patients

To gain better insights into the therapeutic potentials of the interferon signature response in cancer patients, we analyzed their association with the five-year survival of 462 patients with cutaneous melanoma (project ID TCGA-SKCM). Thirty-six genes (77% of all genes examined) that are well-known cancer-associated genes or have been verified in this study were associated with either significantly favorable or poorer (*P*<0.05) five-year survival rates (**Fig. 7a**, **Supplementary Fig. 7** and **Supplementary table 1**). In particular, eighteen interferon response genes (e.g. *Stat1*, *Irf1*), twelve T cell activation and antigen presentation-related genes (e.g. *Cxcl9*, *Cxcl10*, *HLA-DOA* (HLA class II histocompatibility antigen, DO alpha chain)) and three pro-apoptosis genes (*Cp*, *Tnfsf10* and *Fas*) showed positive associations with favorable survival rates (**Fig. 7a**). These findings further corroborated the therapeutic importance of the interferon signature response in melanoma patients as validated in the murine melanoma model. On the other hand, the M2 TAM phenotype gene, *Arg1*, and two pro-angiogenesis/tumor genes (*Cxcl14* and *Mmp13* (matrix metallopeptidase 13)) were associated with poorer survival (**Fig. 7a**). As TAMs are known to display an M2-like phenotype with increased *Arg1* expression and angiogenesis-promoting effects in multiple cancers (4, 5), the above findings thus provide new evidence for TAM-based prognosis. We next examined potential associations with expression profiles of the abovementioned genes and overall survival of cutaneous melanoma patients. In patients with shorter than 321 days (n=50) or longer than 4407 days (n=50), the expression profiles of all thirty-six genes were consistent with their prognosis values. In particular, patients with the longest overall survival harbored higher expression of genes involved in the interferon signature response, T cell activation and antigen presentation, apoptosis, but not the angiogenesis (**Fig. 7b**). Collectively, these findings provide fresh perspectives on the importance of interferon signature response as a favorable prognosis marker, at least in melanoma patients.

**Figure 7.**
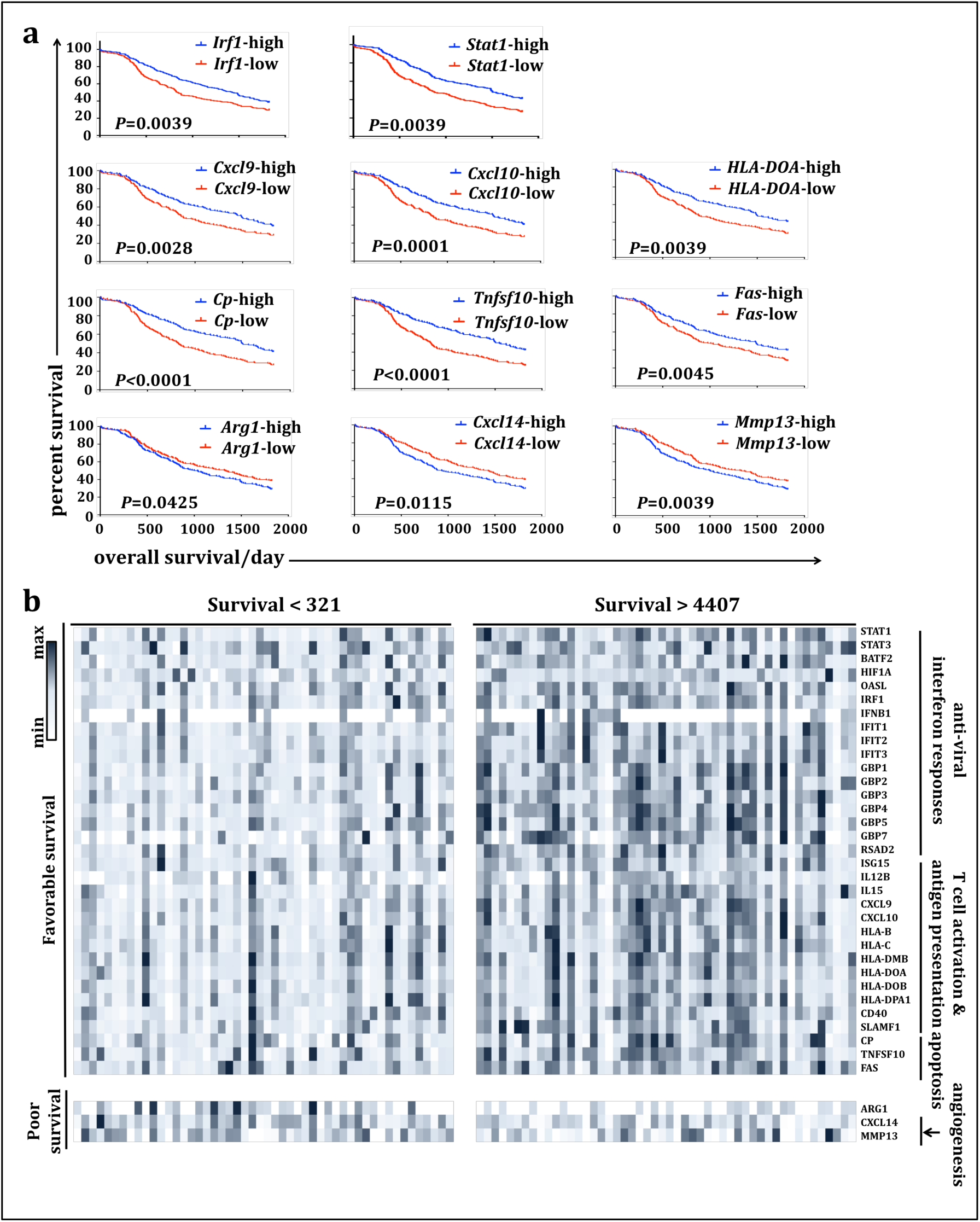
Interferon signature response genes are favorable prognosis markers in cutaneous melanoma. (**a**) Kaplan-Meier curves showing representatives of thirty-six genes with significant associations (*P*<0.05; log-rank test) between their expression levels and patient survival. Gene expression levels were scored as above (high) or below (low) the median expression, before examining their associations with the five-year survival of cutaneous melanoma patients (n=462; project ID TCGA-SKCM) (36). (**b**) Heat map showing mRNA expression of thirty-six genes identified in (**a**). Data presented are from individuals with overall survival shorter than 321 days (n=50) or longer than 4407 days (n=50). See also **Supplementary Fig. 7**.

## Discussion

Instead of recombinant interferons (7-11) and pDC-based pathogen recognition receptor agonists (12-15), we provide here, an alternative and novel strategy that targets TAMs to mount the anti-tumor interferon responses. During viral infection, pDC has long been considered as the major source of type I interferons upon TLR7 and TLR9 ligation (30, 43), whereas macrophages are purportedly activated by TLR3 and TLR4 agonists to produce type I interferons (30). Surprisingly during TLR-TLR crosstalk, TLR3- and TLR4- induced interferon responses in macrophages are constrained by the activated TLR7. As aberrant TLR7 signaling is known to cause autoimmune disorders (44, 45), it is conceivable that macrophages are programmed to prevent excessive production of type I interferons as earlier implied (22). Indeed, interferon response genes became detectable in TLR7-activated macrophages only when the MEK1/2 pathway was inhibited. This is in agreement with previous reports that MEK1/2 pathway is activated by the Tpl2 MAP3K (46), a kinase that downregulates TLR4- and TLR9- induced IFN-β production in macrophages but not pDCs (22). The suppression on IRF1-mediated interferon signature response is thus another fundamental physiological function of the MEK1/2 pathway, which may be therapeutically exploited as an innovative strategy to mount anti-tumor interferon responses.

Despite the above roles in constraining interferon production, the MEK1/2 pathway is also essential for the proliferation and survival in many cell types including macrophages (13, 38). Consistently, we found that inhibition of MEK1/2 pathway dampened macrophage proliferation and increased cell death *in vitro*. However, similar to previous report in CT26 tumors (39), the granzyme B^+^ CD8^+^ T cells were not reduced by monotherapy with MEK1/2 inhibitor *in vivo*. The numbers of neutrophils and Foxp3^+^ Treg cells were also unaffected by the MEK1/2 inhibitor, suggesting that the severe side effects caused by MEK1/2 inhibitors are dose-dependent and cancer-specific (2, 13, 21). Furthermore, MEK1/2 inhibitor-based immunotherapies completely depend on the *RAS*/*RAF* mutations carried by the heterogeneous tumor cell populations (2, 39, 41). This selective effect explains our observation that MEK1/2 inhibitor did not alter the progression of non-mutated B16F10 tumors. Although high doses of MEK1/2 inhibitors caused severe side effects (2, 21), combination therapies using tolerable doses of MEK1/2 inhibitors with BRAF inhibitor (13) or immune checkpoint blockade (39) generated appealing outcomes. Similar outcomes were also observed during our combination therapy *in vivo*. One explanation may be the relatively lower resistance during combination therapies. Another reason could be that MEK1/2 inhibitor caused tumor regression by targeting both the *RAS*/*RAF*-dependent cancer cells and the *Tpl2*-dependent TAMs (21). Hence, our unique combination therapy actually enables a concurrent targeting of the tumor-associated immune cells like TAMs, regardless of the *RAS*/*RAF* mutations in cancer cells.

The immunogenic potentials of TLR7 agonist-based adjuvants are known to be mediated by type I interferon signaling via the pDCs (14, 29), which subsequently mount anti-tumor responses via CTLs and NK cell, or via the constraint of suppressive Treg cells. Although monotherapy with TLR7 agonist significantly improved survival of tumor-bearing mice, the anti-tumor effects were significantly enhanced when combined with the MEK1/2 inhibitor. Therefore, it is likely that the relatively limited efficacies of TLR7 agonists *in vivo* may be due to its incompetency in activating TAMs, which normally display an M2-like phenotype by secreting immunosuppressive cytokines like IL-10 (4, 5, 29, 47). Indeed, we found that *in vitro* TLR7 stimulation led to mixed phenotypes in TAMs that were further reprogrammed to MHC II^+^ CD163^-^ M1-like TAMs when combined with the MEK1/2 inhibitor. Similar results were found regarding the expression of well-known M2 and M1 phenotype markers. On the other hand, the counter-balancing effects of IL-10 signaling were abolished during the combination treatment. As a result, we observed significantly improved mice survival and elevated expression of anti-tumor chemokines (*Cxcl9* and *Cxcl10*) (48), and pro-apoptosis genes (*Tnfsf10/Trail* and *Fas*) (49) following combination therapy. Although numerous studies and clinical trials have tested how elimination of TAM populations and prevention of macrophages recruitment may affect the therapeutic outcomes (4, 5), these strategies somehow ignored the plasticity and anti-tumor potentials of TAMs. Intriguingly, our studies proved that introducing MEK1/2 inhibitor into TLR7 agonist monotherapy fully unlocked the anti-tumor efficacies of TAMs while subverting their immunosuppressive effects.

In summary, the superior efficacies of our combination therapy are at least four-fold (**Supplementary Fig. 8**): (1) elimination of the immunosuppressive roles of TAMs, (2) exploitation of the anti-tumor type I interferons, cytokines and pro-apoptotic molecules from reprogrammed immunostimulatory TAMs, (3) maintenance of the anti-tumor effects of TLR7-based adjuvant mainly via pDCs, and (4) maintenance of the antineoplastic effects of MEK1/2 inhibitor mainly in *RAS*/*RAF*-mutated cancers. Future immunotherapies may be improved by adopting our novel strategy of combination therapy together with the immune checkpoint blockades in the *RAS/RAF*-mutated cancers.

## Methods

### Animals

Age- and sex-matched mice were used in all experiments. WT C57BL/6JInv were purchased from InVivos Pte Ltd. B6.129S2-*Irf1^tm1Mak^*/J (*Irf1*^-/-^) and B6.129S(Cg)-*Stat1^tm1Dlv^*/J (*Stat1*^-/-^) were from the Jackson Laboratories and backcrossed to C57BL/6J backgrounds for over ten generations. Femurs and tibias of B6.129P2(SJL)-*Myd88^tm1.1Defr^*/J (*Myd88*^-/-^) were kindly provided by Dr. Norman Pavelka (Singapore Immunology Network, SIgN, Singapore). *Stat1*^ind^ mice were generously provided by Professor Mathias Muller (Inst. Of Animal Breeding and Genetics, University of Veterinary Medicine, Vienna). Animals were maintained at the Department of Comparative Medicine, National University of Singapore.

### Cell culture

BMDM were derived from the bone marrow cells (35). Briefly, femurs and tibias were extracted and flushed using phosphate-buffered saline (PBS; 137 mM NaCl, 2.7 mM KCl, 8 mM Na_2_HPO_4_, 2 mM KH_2_PO_4_, pH 7.4). Red blood cells were then lysed using ACK lysing buffer (Gibco, Thermo Fisher Scientific) and remaining cells were filtered through 70 μm cell strainers (Miltenyi Biotec) before plating at a density of 1.5 x 10^6^ cells/ml. Culture medium were Dulbecco’s modified Eagle’s medium (DMEM; Gibco, Thermo Fisher Scientific) supplemented with 10% (v/v) fetal bovine serum (FBS; HyClone, Thermo Fisher Scientific), recombinant macrophage colony-stimulating factor (100 U/ml; ebioscience, Thermo Fisher Scientific) and Pen Strep (1:100 dilution; Gibco, Thermo Fisher Scientific). Non-adherent cells were then removed from downstream applications on day seven. Other cells including the J774.1 macrophage cell line (ATCC) and B16F10 melanoma cells (ATCC^®^ CRL-6475™) were all cultured in DMEM supplemented with 10% (v/v) FBS and Pen Strep (1:100 dilution). BMDM and J774.1 cells were plated at a density of 1 × 10^6^ cells/ml and 0.4 × 10^6^ cells/ml, respectively. All cells were cultured in an incubator at 37°C with 5% CO_2_.

### Reagents and antibodies

Poly(I:C) (LMW; Invivogen) was used at 10 μg/ml. R848 (Resiquimod; Invivogen) was used at 25 ng/ml for *in vitro* studies and 3 mg/kg for *in vivo* melanoma model. Pam3CSK4 (Calbiochem, EMD Biochemicals) was used at 10 ng/ml. *Escherichia coli* 055:B55 LPS (Sigma-Aldrich) was used at 100 ng/ml. Recombinant mouse IL-10 (cat. no. 575802), IFN-β1 (cat. no. 581302) and IFN-γ (cat. no. 575302) proteins were purchased from Biolegend.

MEK1/2 inhibitors, including MEKi-U (U0126; cat. no. #9903; Cell Signaling Technology), MEKi-A (AZD6244, selumetinib; Selleck, USA) and MEKi-P (PD0325901; Selleck, USA) were used at 10 μM, 0.5 μM and 0.5 μM, respectively. mRNA synthesis was inhibited with 5 μg/ml actinomycin D (Sigma-Aldrich). Inhibitors targeting p38 MAPK (p38i, SB203580; cat. no. #5633) and JNK MAPK (JNKi, SP600125; cat. no. #8177) were purchased from Cell Signaling Technology and used at 10 μM final concentrations. NF-κB inhibitor (NF-κBi, BAY 11-7085; sc-202490) were from Santa Cruz Biotechnology. All inhibitors, except for actinomycin D, were added one hour prior to the indicated stimulations (24, 25).

Antibodies used in this study include anti-IRF1 (sc-640), anti-GAPDH (sc-32233), anti-STAT1 (sc-271661), anti-IκBα (sc-1643) and anti-Cot/Tpl2 (sc-720) from Santa Cruz Biotechnology; anti-p65/RelA (ab-7970), anti-STAT3 (ab-119352) and anti-pSTAT3 Y705(ab-76315) from Abcam; anti-pSTAT1 S727 (#9177), anti-pSTAT1 Y701 (#9171), anti-p44/42 MAPK (ERK1/2; #9107s), anti-pERK1/2 T202/Y204 (#9101s), anti-p38 MAPK (#9212s), anti-p-p38 T180/Y182 (#9211s) and anti-p-p65/RelA S536 (#3033P) from Cell Signaling Technology; IgG isotype control, anti-mouse CD210 (IL-10R) and anti-mouse CD119 (IFNGR) from ebioscience, Thermo Fischer Scientific; and anti-mouse IFNAR (cat no. 127323) from Biolegend. All antibodies were used at 1:1000 dilution except for those used for the neutralization assays, including IgG isotype control (10 μg/ml unless otherwise specified), anti-IL-10R (10 μg/ml unless otherwise specified), anti-IFNAR (10 μg/ml) and anti-IFNGR (10 μg/ml). For neutralization studies, isotype controls and all antibodies were added one hour before the indicated stimulations.

For reconstitution of STAT1 in *Stat1*^ind^ mice, harvested BMDM were incubated with doxycycline (cat. no. 17086-28-1; Sigma-Aldrich) at indicated concentrations for 24 hours (35). Culture medium was then removed and replenished with fresh culture medium before downstream analysis.

### Murine melanoma model

Mycoplasma-free B16F10 (ATCC^®^ CRL-6475™) cells were harvested during logarithmic phase and resuspended in sterile PBS. An inoculum of 0.2 million B16F10 cells in 100 μl PBS was injected subcutaneously (s.c.) into the left flank of C57BL/6 mice. Seven days after injection, mice with palpable tumors were randomly divided into different groups and treated as indicated (**Fig. 6A**). TLR7 agonist R848 was reconstituted according to the manufacturer’s instructions and administered intratumorally (i.t.) at 3 mg/kg every two days. MEK1/2 inhibitor MEKi-A (AZD6244) was formulated in sterile vehicle with 0.5% (Hydroxypropyl)methyl cellulose (Sigma-Aldrich) and 0.1% polysorbate 80 (v/v) (Sigma-Aldrich) as described before (34). In total, 50 mg/kg MEKi-A was administered twice daily via oral gavage (p.o.). Accordingly, control group was treated with vehicle twice daily (B.I.D.) and sterile PBS at 3 ml/kg every two days. Tumor size was monitored as an area (longest dimension × perpendicular dimension) using a digital caliper thrice per week until the experimental endpoints (tumor size > 100 cm^2^) as reported before (50).

### Interferon signature response in the Cancer Genome Atlas (TCGA) data

Associations between expression levels of twenty-one anti-viral interferon response-related genes, twenty-four T cell activation and antigen presentation-related genes, five pro-apoptosis genes, four macrophage-related genes, three pro-angiogenesis genes and five-year survival patient survival were analyzed. Clinical data for 462 TCGA skin cutaneous melanoma (SKCM) patients (51) were downloaded from the TCGA data portal using UCSC Xena (http://xena.ucsc.edu/), including the overall survival in days and gene expression data from Illumia HTseq 2000 RNAseq platform. Data used were all listed in the **Supplementary table 1**. The median follow-up from diagnosis recorded was 1124 days with a range of 6 to 11252 days. For each gene, we used a similar method as reported before (36) to score the gene expression as above or below the median expression, and examined their associations with truncated five-year survival (1825 days) using log-rank tests. Genes with higher or lower expression profiles associated with favorable survival (*P*<0.05) were considered as prognosis markers for favorable or poor survival, respectively. In total 36 genes were identified with 33 favorable and 3 poor prognosis markers.

### Flow cytometry staining and acquisition

Single-cell suspensions were prepared and stained with fluorescence conjugated antibodies at a 1:200 dilution. Cells were stained with fixable viability dye eFluor™ 506 (ebioscience, Thermo Fisher Scientific), blocked with 2.5 μg/ml anti-CD16/32 (clone 93; ebioscience, Thermo Fisher Scientific) and then stained for flow cytometry with the following antibodies: CD45-eF450 (104), CD11b-FITC (M1/70), MHC II-PE-Cy7 (M5/114.15.2), F4/80-PE-Cy5 (BM8), F4/80-biotin (BM8), Ki67-PerCP-eF710 (SolA15), CD3ε-biotin (145-2C11), CD4-APC-Cy7 (GK1.5), CD4-PE (GK1.5), CD8α-PerCP-eF710 (53-6.7), CD19-biotin (1D3), NK1.1-APC (PK136), CD80/B7.1-PE (16-10A1), granzyme B-PerCP-eF710 (NGZB) and Foxp3-PE-Cy7 (fjk-16s) from ebioscience, Thermo Fisher Scientific; Ly6G-BV605 (1A8), streptavidin-BV605, CD8α-PE (53-6.7), CD68-PE (FA-11), CD206-PE (MR6F3) and CD163-PE (TNKUPJ) from Biolegend and CD64-APC (REA286) from Miltenyi Biotec. Following staining with the cell surface markers, cells were fixed and permeabilized using ebioscience™ Foxp3/Transcription Factor Staining Buffer Set (ebioscience, Thermo Fisher Scientific) following the manufacturer’s instructions. For cell proliferation and cell death analysis, single-cell suspensions were stained with fixable viability dye eFluor™ 450 (ebioscience, Thermo Fisher Scientific) followed by intracellular staining of Ki67-PerCP-eF710. All samples were filtered through 60 μm cell strainers before analysis using BD LSRFortessa™ (BD Biosciences) and FlowJo software (FlowJo, LLC.).

### Quantitative RT-PCR

Total RNA was purified using TRIzol™ reagent (Invitrogen, Thermo Fisher Scientific) and reverse transcribed using SuperScript III First-Strand Synthesis System (Thermo Fisher Scientific) according to the manufacturer’s instructions. Complement DNA (cDNA) templates were then analyzed using GoTaq^®^ qPCR Master Mix (Promega Corporation) on a LightCycler^®^ 480 Instrument II (Riche Life Science). The abundances of all genes tested were normalized to the housekeeping gene hypoxanthine phosphoribosyltransferase 1 (*Hprt*). The primers used in this study were all listed in **Supplementary table 2**.

### Western blotting

Cell lysate was prepared using radioimmunoprecipitation assay buffer (1 x RIPA buffer; 50 mM Tris pH7.5, 0.5% deoxycholate, 150 mM NaCl, 1% NP-40, 0.1% SDS, 1mM EDTA) supplemented with protease inhibitor cocktail (Sigma-Aldrich) following the manufacturer’s instructions. Cell lysate was then processed for SDS-PAGE, followed by blocking with 5% skim milk (Sigma-Aldrich) and incubation with indicated primary antibodies. The blots were then incubated with horseradish peroxidase–conjugated rabbit or mouse secondary antibodies (Sigma-Aldrich) and visualized using WesternBright ECL (Advansta) in an ImageQuant LAS 4000 mini system (GE Healthcare). The blots for loading controls were reprobed after stripping using Restore™ Western Blot Stripping Buffer (Thermo Fisher Scientific) following the manufacturer’s instructions.

### ELISA

IL-10 secretion was determined using the mouse IL-10 ELISA set (BD Biosciences Inc) according to the manufacturer’s instructions.

### RNA purification for GeneChip

RNA isolation from stimulated BMDM was performed using TRIzol™ reagent (Invitrogen, Thermo Fisher Scientific) and RNeasy^®^ Plus Mini Kit (Qiagen) according to the manufacturer’s instructions. Genomic DNA was removed using Ambion™ DNase I (RNase-free) (Thermo Fisher Scientific) before elution. Quantities and qualities of purified RNA were assessed using NanoDrop Spectrophotometer (Thermo Fisher Scientific) and Agilent Bioanalyzer (Agilent Technology). Samples with no significant contamination and with RNA integrity numbers (RIN) larger than 8.0 were included for downstream analysis.

### Whole transcriptome analysis

Double stranded complementary DNA (cDNA) was generated, fragmented and end-labeled using the GeneChip^®^ WT PLUS Reagent Kit (Thermo Fisher Scientific). Labeled cDNA was then processed and hybridized to GeneChip^®^ Mouse Transcriptome Array 1.0 (Affymetrix, Thermo Fisher Scientific) using AFX Fluidics 450 Station (Thermo Fisher Scientific) following the manufacturer’s instructions. GeneChips were scanned on a GC3000 G7 Scanner (Thermo Fisher Scientific). Raw microarray files were then extracted, annotated and normalized by the recommended RMA algorithm of Partek Genomic Suite (v6.6) (Partek Incorporated) with the core meta-probe set. Probe set information was then log2 transformed and consolidated at gene levels. Analysis of variance (ANOVA) was performed with contrasts set against corresponding non-treated (NT) controls. Genes differentially expressed by no less than 1.5 folds with adjusted *P* values<0.05 unless otherwise specified were shortlisted for downstream analysis.

### Gene set enrichment analysis

Shortlisted genes were imported for the PANTHER overrepresentation test (released on 20171205; Gene Ontology Consortium)using the default Fisher’s Exact Test with false-discovery rate (FDR) correction for multiple tests (52, 53). Gene lists with FDR < 0.05, raw *P* value < 10^-4^ and fold enrichment > 10 unless otherwise specified were considered as significantly enriched gene sets.

### STRING PPI test

Differentially expressed genes were analyzed using STRING database (version 10.5) with default settings and minimum required interaction score (0.400) (54). The active interaction sources were customized to include only known interactions from curated databases and experimental data.

### Statistics

Data are presented as means ± S.D. unless otherwise specified. Statistical analysis was performed using Prism 6.0 software (Graph Pad software). Unpaired Welch’s t-test, log-rank test, or analysis of variance (ANOVA) with Bonferroni’s or Tukey’s correction for multiple comparisons were used to determine the statistical significances. *P* value<0.05 were considered significant. Researchers were not blinded to the experimental groups and all samples were included for the analysis.

### Data availability

Data used to support the findings of this study are presented in figures or supplementary materials accompanying the main text or are available from the corresponding author upon request. Whole transcriptome data are available at the Gene Expression Omnibus (GEO) repository with GEO accession number GSE123512.

### Study Approval

All experiments were carried out in accordance with institutional guidelines prescribed by the Institutional Animal Care and Use Committee, National Univesity of Singapore (protocol number R13-5667 and R17-1205).

## Author contributions

L. Yang designed and performed all experiments. J.L. Ding conceived and supported all studies. L. Yang and J.L. Ding analyzed the data and wrote the manuscript.

## Acknowledgements

We thank the National Medical Research Council (NMRC/CBRG/0055/2013) and MOE (R-154-000-A76-114) for financial support. We are grateful to Professor Mathias Muller (Inst. Of Animal Breeding and Genetics, University of Veterinary Medicine, Vienna) for providing *Stat1*^ind^ mice. Femurs and tibias from *Myd88^-/-^* mice were kindly provided by Dr. Norman Pavelka (Singapore Immunology Network, SIgN, Singapore). We thank (Mr.) Er Junzhi for his comments on our manuscript. We are grateful to Dr. Ng Lai Guan (SIgN) for technical advice. L. Yang is supported by the NUS Research Scholarship, Faculty of Science, National University of Singapore.

## Supplementary figures

**Supplementary Fig. 1.**
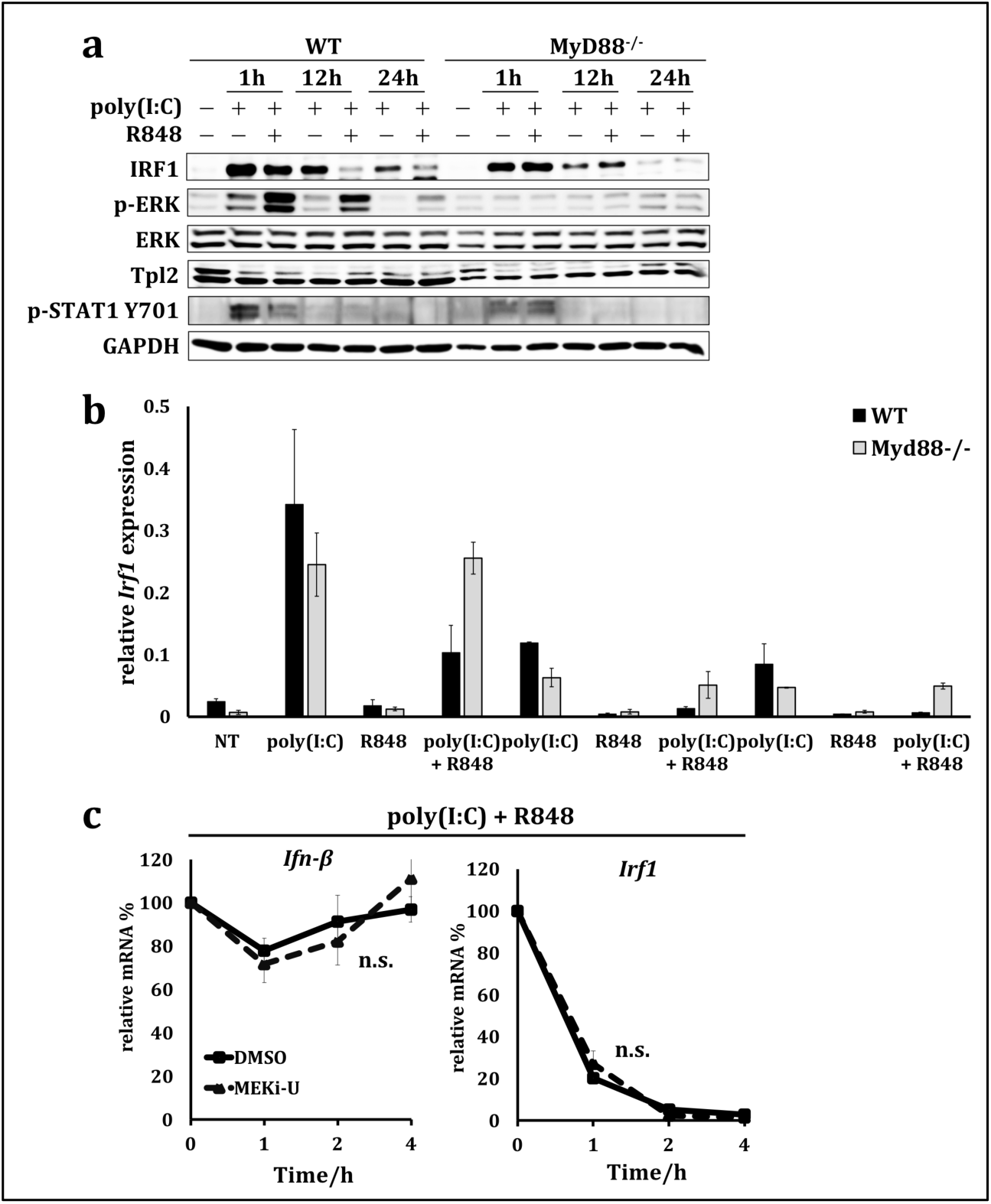
TLR7 agonist activates MyD88-MEK1/2-mediated suppression in macrophages. (**a**) Immunoblot analysis of IRF1, total and p-ERK, Tpl2/Cot and p-STAT1 Y701. Primary macrophages from either *Myd88*^-/-^ or WT mice were treated with TLR7 agonist R848 and/or TLR3 agonist poly(I:C) as indicated. (**b**) mRNA analysis of *Irf1* in BMDM cells treated similarly as in (**a**). (**c**) mRNA stabilities of *Irf1* and *Ifn-β*. J774.1 macrophage cells were treated with both R848 and poly(I:C) in the presence or absence of MEK1/2 inhibitor MEKi-U. At four hours post stimulation, cells were assayed as zero-time point. *De novo* mRNA synthesis was then inhibited using actinomycin D for indicated time durations. Gene expression was all normalized to *hprt* and presented as percentages of the mRNA levels at zero-time point. n.s, not significant. Data are presented as mean ± S.D. or one representative of two (**a**-**b**) and three (**c**) independent experiments.

**Supplementary Figure 2.**
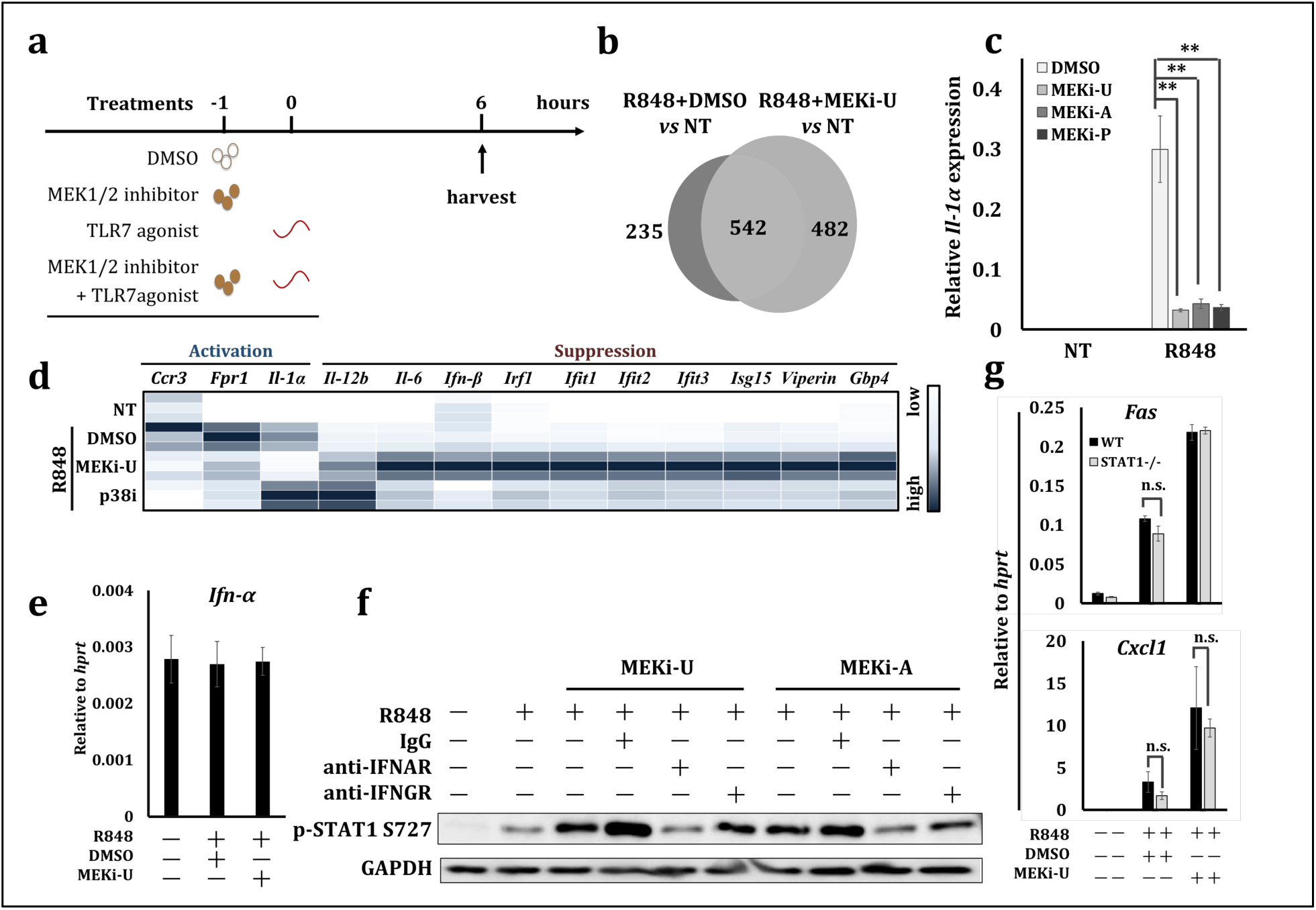
MEK1/2 inhibitor synergizes with TLR7 agonist in macrophages. (**a**) treatment scheme for Fig. 2a. (**b**) Venn diagram showing numbers of DEGs from the whole transcriptome analysis. (**c**) mRNA expression of *Ifn-α*. BMDM were stimulated for six hours in the presence or absence of MEK1/2 inhibitors as indicated. (**d**) Heat map showing mRNA expression of genes activated or suppressed by the MEK1/2 pathway or p38 MAPK pathway. BMDM were activated via R848 for 6 hours in the presence or absence of MEKi-U and p38 inhibitor p38i. Scale bars are presented as a range from low to high expression relative to each gene. (**e**) mRNA expression of proinflammatory gene *Il-1α*. BMDM were stimulated similarly as indicated in (c). (**f**) Immunoblot analysis of p-STAT1 S727. BMDM were stimulated with TLR7 agonist R848 for twelve hours in the presence or absence of MEK1/2 inhibitors, IgG isotype control, anti-IFNAR and anti-IFNGR antibodies. (**g**) mRNA expression of pro-apoptotic gene *Fas* and chemokine *Cxcl1*. Cells were treated similarly as in (**e**). *0.01<*P*<0.05, **0.001<*P*<0.01 (unpaired Welch’s t-test or two-way ANOVA with Tukey’s correction). n.s., not significant. Data are presented as relative expression (**d**), mean ± S.D. (**c**, **e**, **g**) or one representative of three (**f**) or four (**c**, **e**, **g**) independent experiments.

**Supplementary Figure 3.**
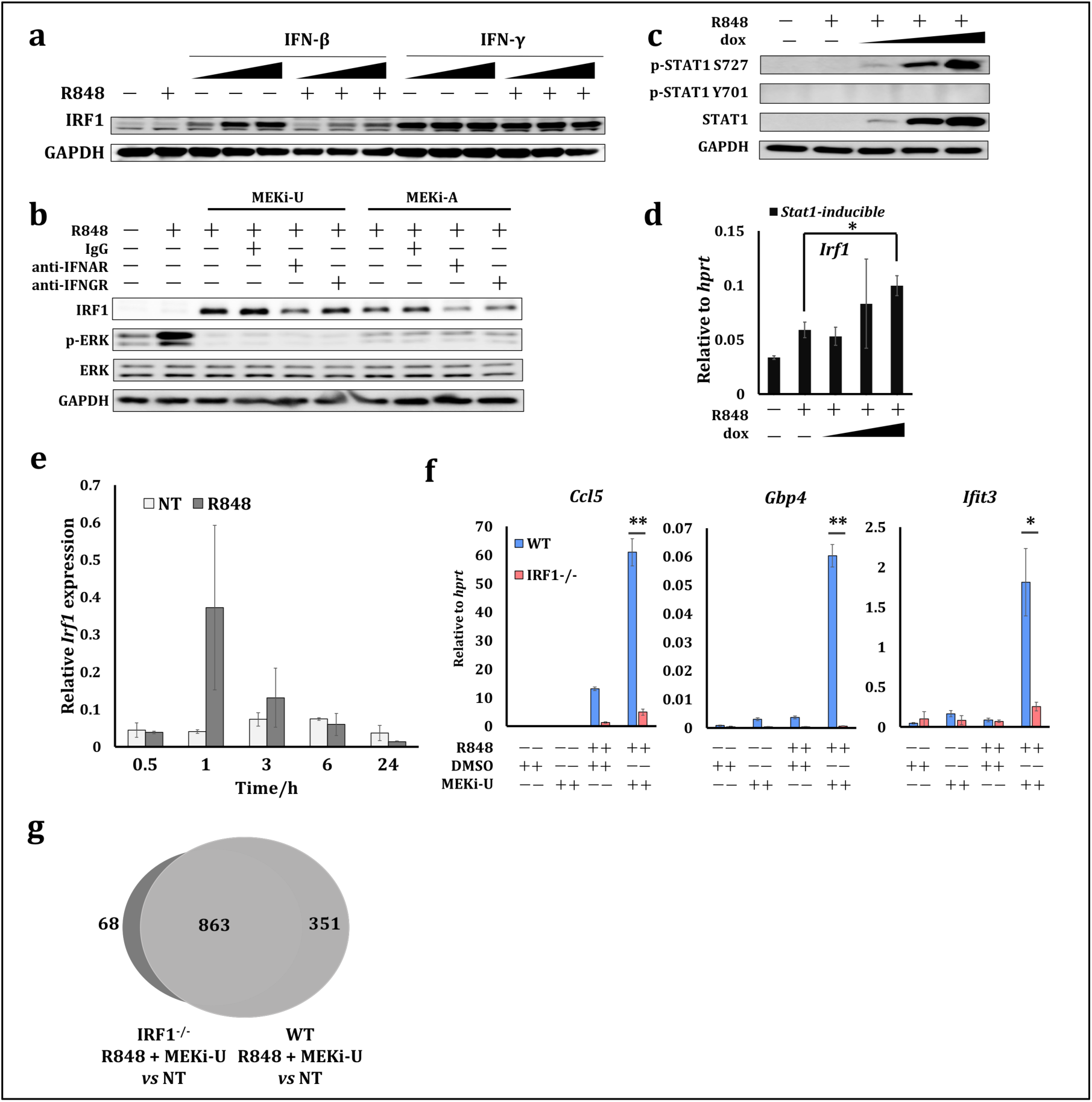
IRF1 is involved in MEK1/2 inhibition-generated interferon signature response. (**a**) Immunoblot analysis of IRF1. J774.1A macrophage cells were treated with TLR7 agonist R848 alone, or together with murine recombinant IFN-β (2 U/ml, 200 U/ml and 2000 U/ml) and IFN-γ (2 U/ml, 200 U/ml and 2000 U/ml) for twelve hours as indicated. (**b**) Immunoblot analysis of IRF1, total and p-ERK. BMDM were stimulated with R848 for twelve hours in the presence or absence of MEK1/2 inhibitors, IgG isotype control, anti-IFNAR and anti-IFNGR antibodies. (**c**) Immunoblot analysis of total STAT1, p-STAT1 S727 and p-STAT1 Y701. BMDM generated from *Stat1^ind^* mice were pre-treated with increasing concentration of doxycycline (0.03125, 0.125 and 0.5 μg/ml) for twenty-four hours (35). Culture medium was then replaced and cells were stimulated with R848 for eight hours. (**d**) mRNA expression of *Irf1*. Cells were similarly treated as in (**c**). (**e**) mRNA expression of *Irf1*. BMDM were stimulated with R848 for indicated time durations. (**f**) Expression of chemokine *Ccl5* and two interferon response genes *Gbp4* and *Ifit3*. BMDM from both WT and Irf1^-/-^ mice were treated with R848 for six hours in the presence or absence of MEKi-U. (**g**) Venn diagram showing numbers of DEGs from whole transcriptome analysis using both WT and IRF1^-/-^ macrophages. *0.01<*P*<0.05, **0.001<*P*<0.01 (unpaired Welch’s t-test). Data are presented as mean ± S.D. or one representative of three (**a**, **c**-**e**) or four (**b**, **f**) independent experiments.

**Supplementary Figure 4.**
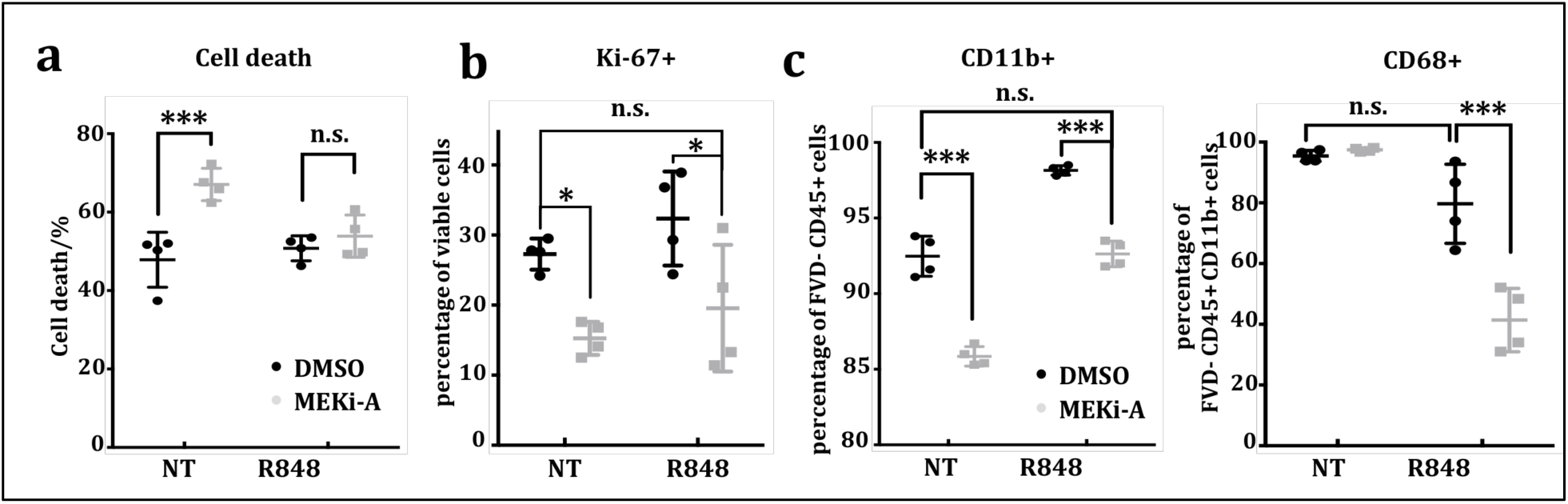
Combination of MEK1/2 inhibitor and TLR7 agonist modulates macrophage survival and proliferation. (**a**, **b**, **c**) Percentages of FVD^+^ dead cells (**a**), FVD^-^Ki-67^+^ proliferating cells (**b**), and CD11b^+^ and CD68^+^ macrophages (**c**) were assessed using flow cytometry. BMDM were assayed similarly as that in (**c-e**). BMDM were stimulated with R848 for twenty-four hours in the presence or absence of MEKi-U. Cells were then harvested and assayed for flow cytometry analysis. *0.01<*P*<0.05, **0.001<*P*<0.01, \*\*\**P*<0.001 (one-way ANOVA with Bonferroni’s correction or two-way ANOVA with Tukey’s correction). n.s., not significant. Data are presented as mean ± S.D. or one representative of two (**a**-**c**, n=4 per group) independent experiments.

**Supplementary Figure 5.**
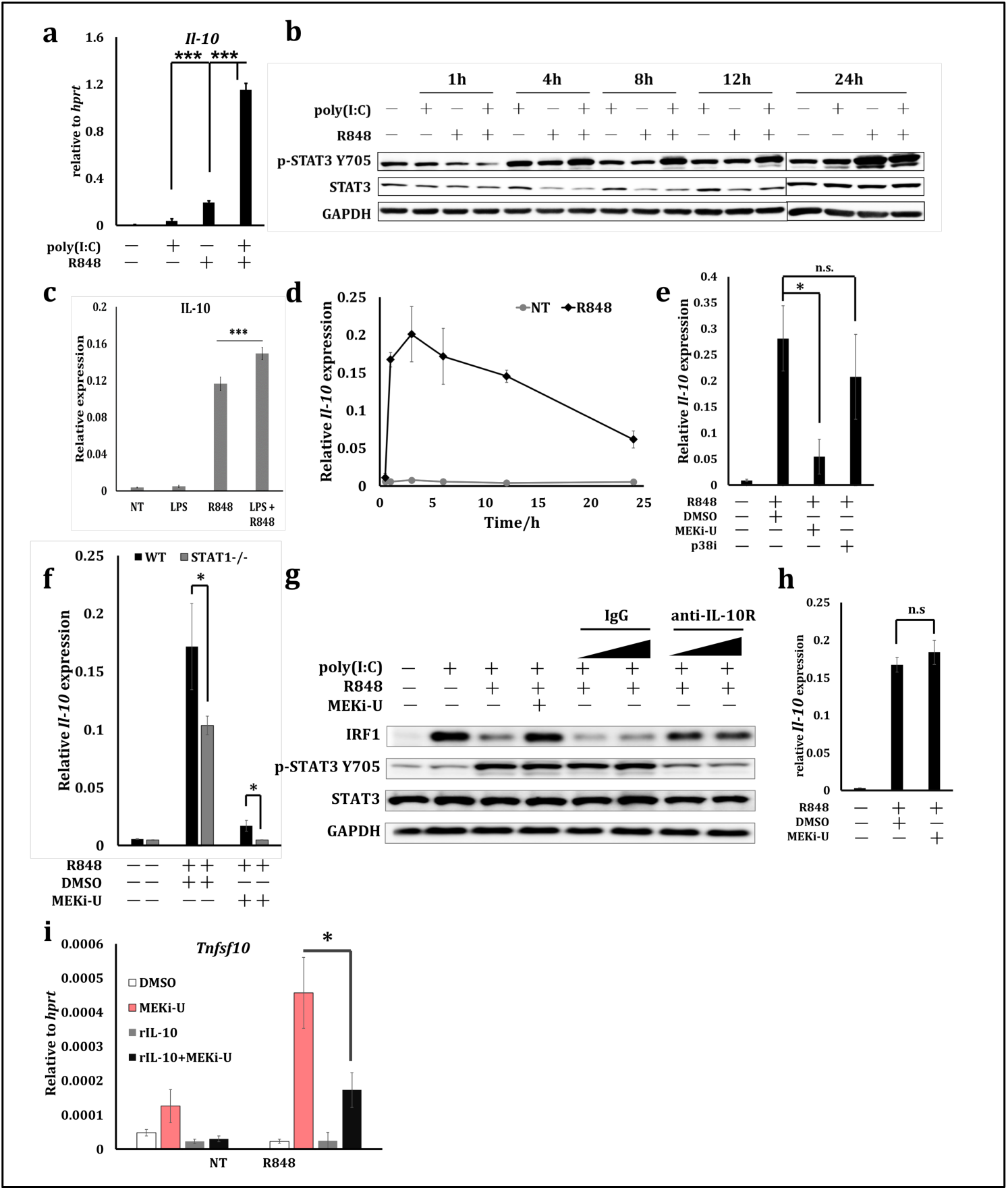
IL-10 signaling suppresses IRF1-mediated interferon signature response. (**a**) mRNA expression of *Il-10*. BMDM were stimulated with single or combination of TLR3 agonist poly(I:C) and TLR7 agonist R848 for twelve hours. (**b**) Immunoblot analysis of total STAT3 and p-STAT3 Y705. J774.1A macrophage cells were stimulated with poly(I:C) and/or R848 for indicted time durations. (**c**) *Il-10* expression in BMDM stimulated with TLR4 agonist LPS and/or R848 for twelve hours. (**d**) mRNA expression of *Il-10* in BMDM stimulated with R848 for indicated time durations. (**e**) mRNA expression of *Il-10* in BMDM stimulated with R848 for six hours in the presence or absence of MEKi-U and p38 MAPK inhibitor p38i. (**f**) *Il-10* expression in BMDM generated from WT and *Stat1*^-/-^ mice. Cells were treated similarly as in (**e**). (**g**) Immunoblot analysis of IRF1, total STAT3 and p-STAT3 Y705. BMDM were stimulated with R848 and poly(I:C) in the presence or absence of MEKi-U, anti-IL-10R antibody (1 μg/ml and 10 μg/ml) and IgG isotype control (1 μg/ml and 10 μg/ml). (**h**) *Il-10* expression in BMDM stimulated with R848 for one hour in the presence or absence of MEKi-U. (**i**) mRNA expression of pro-apoptotic gene *Tnfsf10*. BMDM cells were stimulated with recombinant IL-10 or R848 for six hours in the presence or absence of MEKi-U. *0.01<*P*<0.05, **0.001<*P*<0.01, \*\*\**P*<0.001 (unpaired Welch’s t-test or one-way ANOVA with Bonferroni’s correction). n.s., not significant. Data are presented as mean ± S.D. or one representative of three (**a-e**, **h**) or four (**f**, **g**, **i**) independent experiments.

**Supplementary Figure 6.**
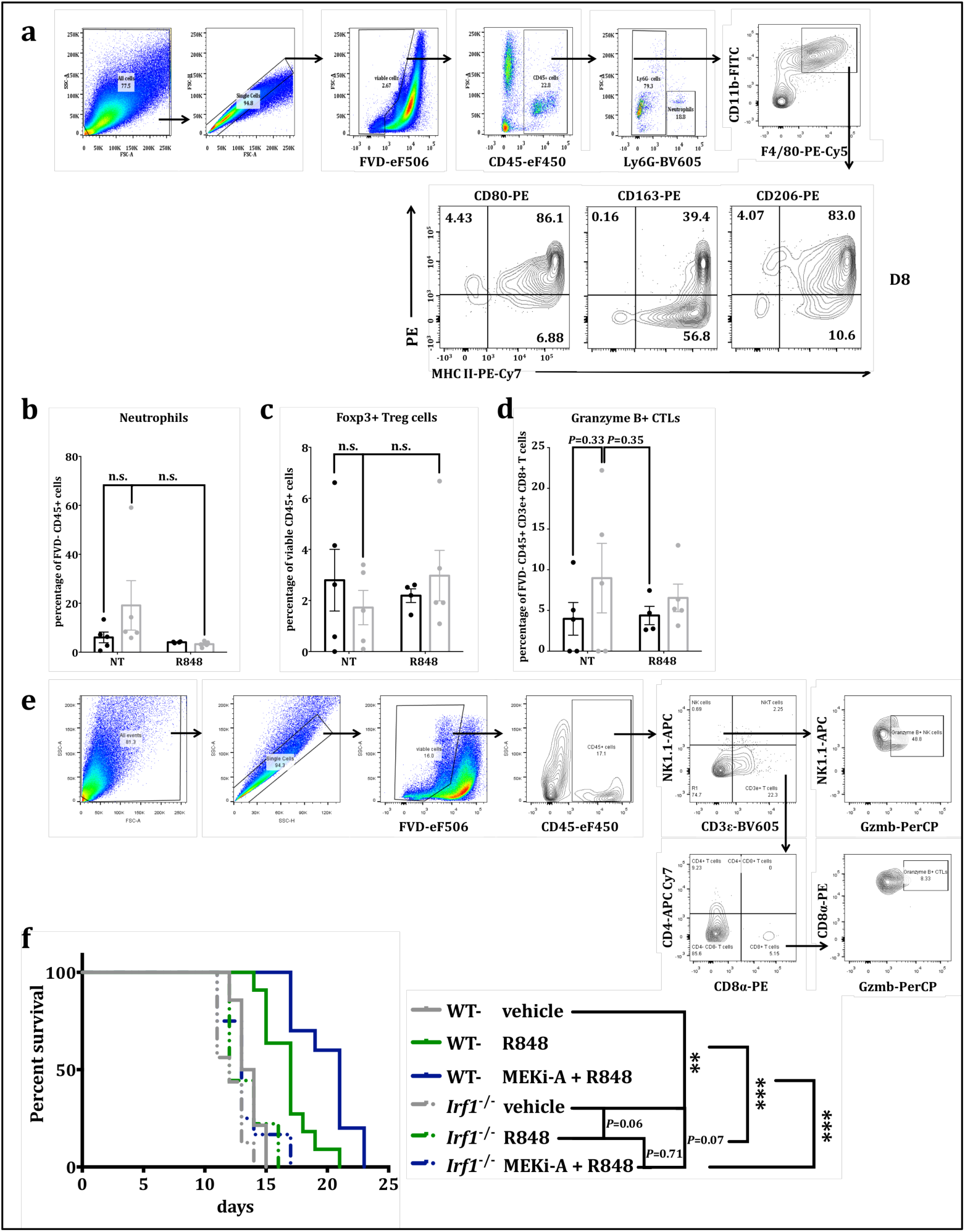
Combination treatment with MEK1/2 inhibitor and TLR7 agonist reduces tumor progression. (**a**) Gating scheme of total TAMs for flow cytometry. B16F10 bearing tumors were allowed to grow for eight days. Tumors were then harvested for single cell suspension preparation and stained for flow cytometry. (**b**, **c**, **d**) Percentages of infiltrating neutrophils (FVD^-^CD45^+^CD11b^+^F4/80^-^Ly6G^+^) (**b**) Foxp3^+^ regulatory T cells (Treg cells; FVD^-^CD45^+^NK1.1^-^CD3ε^+^CD4^+^CD8^-^Foxp3^+^) (**c**) and granzyme B^+^ NK cells (FVD^-^CD45^-^CD3ε^-^NK1.1^+^Granzyme B^+^) (**d**) in B16F10 tumors treated with the same scheme as depicted in Fig. 6a. (**e**) Gating scheme of CTLs and granzyme B^+^ (Gzmb^+^) NK cells for flow cytometry. Tumors were similarly treated as in (**a**). (**f**) Kaplan–Meier survival curves of tumor-bearing WT mice receiving vehicle (n=14), TLR7 agonist R848 (n=11) and combination therapy (MEKi-A and R848, n=10), and of *Irf1^-/-^* mice receiving vehicle (n=16), R848 (n=9) and combination therapy (MEK1/2 inhibitor and R848, n=12). Murine B16F10 subcutaneous melanoma was established and treated similarly as shown in Fig. 6a. *0.01<*P*<0.05, **0.001<*P*<0.01, \*\*\**P*<0.001 (unpaired Welch’s t-test or log-rank test). n.s., not significant. Data are presented as mean ± S.D. or one representative of three (**a**) or two (**b**-**d**, n=4 per group) independent experiments.

**Supplementary Figure 7.**
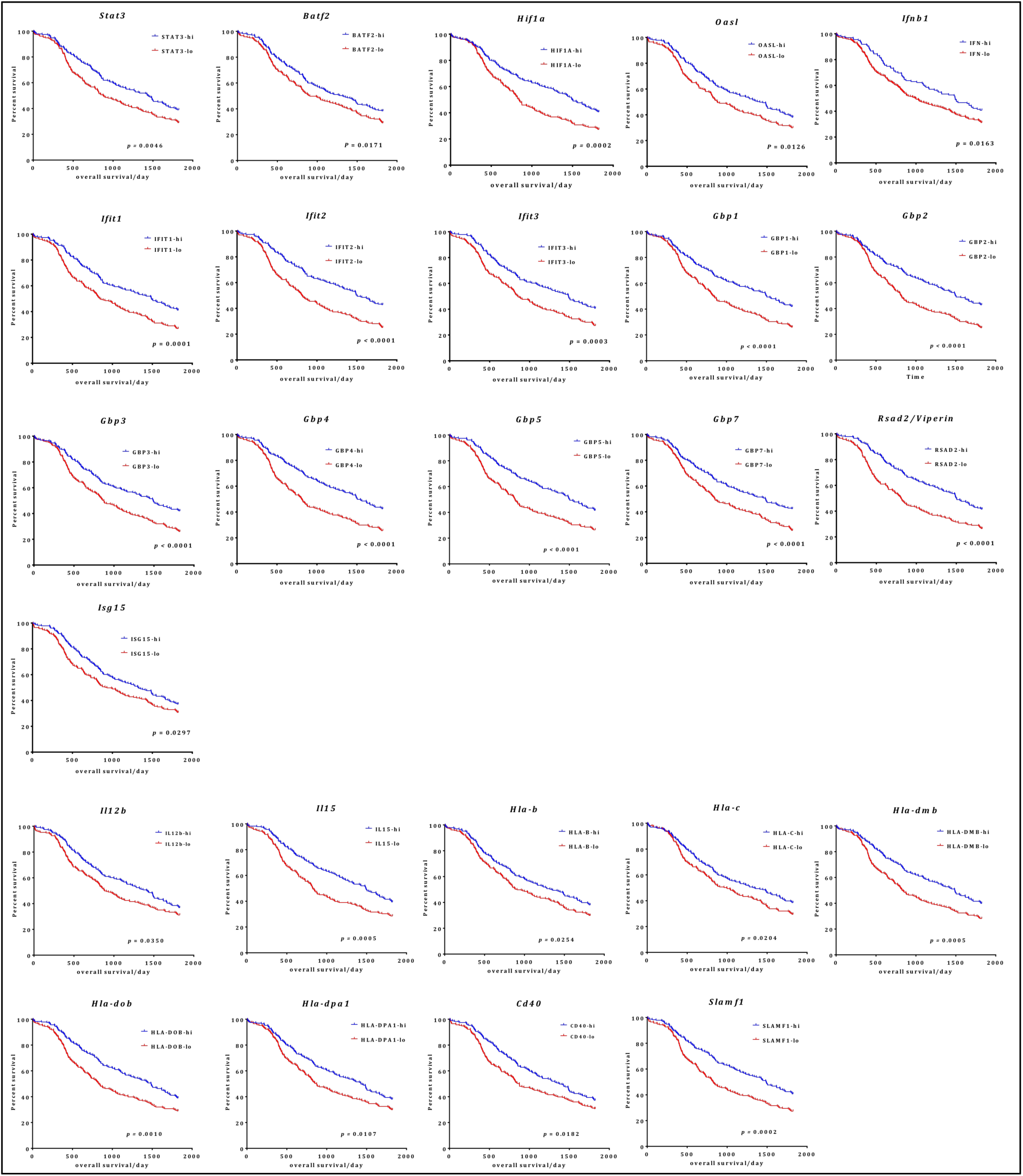
Kaplan-Meier curves showing interferon signature response as favorable prognosis markers. Kaplan-Meier curves showing all genes (except those presented in Fig. 7a) with significant associations (*P*<0.05; log-rank test) between their expression levels and patient survival. Gene expression levels were scored as above (high) or below (low) the median expression, before examining their associations with the five-year survival of cutaneous melanoma patients (36).

**Supplementary Figure 8.**
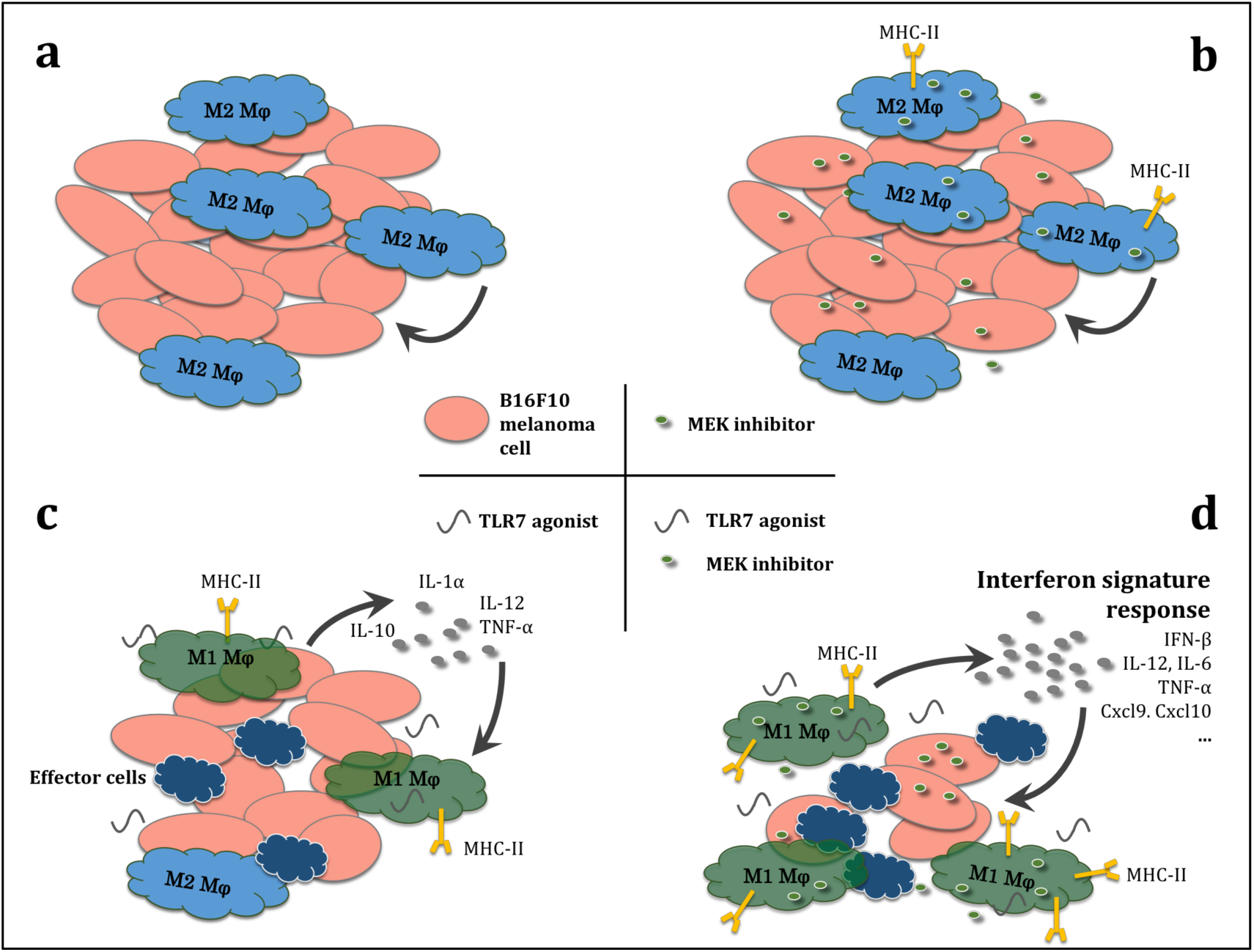
Hypothetical model for the in vivo combination therapy. B16F10 melanoma cell progression and tumor-associated macrophages (TAMs) phenotypes (**blue** for M1 TAM; **green** for M2 TAM) are shown as results of the indicated therapies. Potential effector molecules, such as the interferon signature response, that are known to enhance tumor-killing effects of effectors cells are highlighted. (**a**) Infiltration of immunosuppressive M2-like TAMs can be observed in B16F10 tumors. (**b**) Monotherapy with MEK1/2 inhibitor does not alter tumor progression in *RAS*/*RAF* WT B16F10 tumors. The proliferation and survival of M2-like TAMs are sustained after MEK1/2 inhibitor treatment, in spite of the increase in MHC II expression. (**c**) Monotherapy with TLR7 agonist reduces tumor progression and extends the survival of tumor-bearing mice, possibly via increased antigen-presentation via MHC II, enhanced adaptive immunity via IL-12 and TNF-α, or other mechanisms as reported before (29). However, the TAMs populations display a mixed phenotype with both M1-like and M2-like phenotype markers. Counter-acting IL-10 signaling is also activated through TAMs, a mechanism that may largely compromise TLR7 monotherapy. (**d**) Combination therapy extends survival, even in comparison with the respective monotherapies. Mechanistically, MEK1/2 inhibitor synergizes with TLR7 agonist to unlock an IRF1-mediated “interferon signature response”, with increased expression of chemokines and cytokines known to modulate TIME.

## Supplementary tables

***Supplementary table 1.** Associations between immune-related gene expression and cutaneous melanoma patient survival (see excel file enclosed separately).*

**Supplementary table 2.**
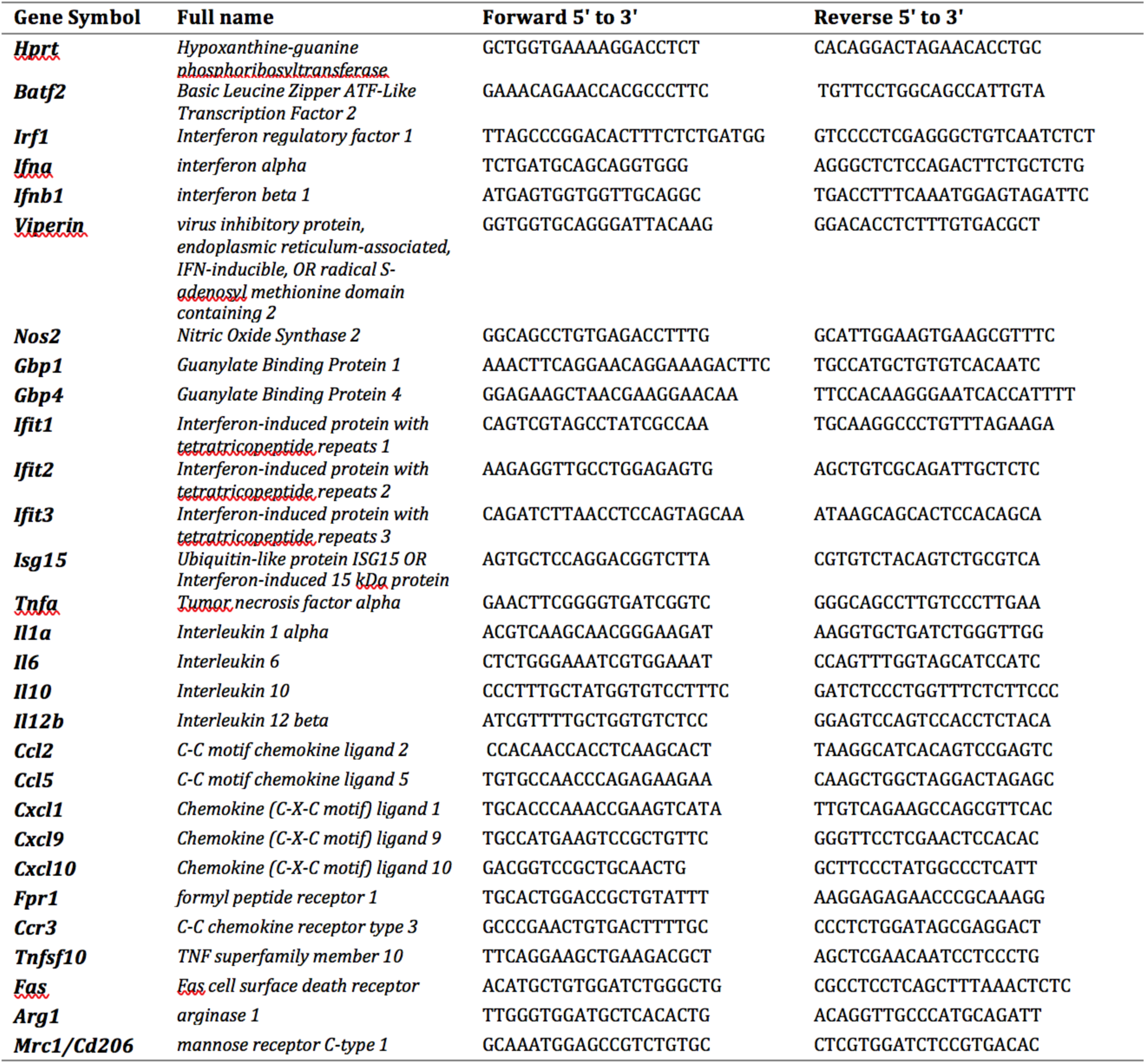
Primers used for quantitative RT-PCR.

